# How are evolutionarily young and old proteins distributed in sequence space?

**DOI:** 10.64898/2026.07.21.739789

**Authors:** Lars A. Eicholt, Agnes Toth-Petroczy, Richard A. Goldstein, Erich Bornberg-Bauer

## Abstract

Protein sequence space is vast due to the combinatorial diversity of 20 amino acids. However, evolution has generated a limited set of “old” canonical protein families sharing evolutionary ancestry, structures and functions. It remains unclear how canonical sequences are placed in sequence space, how recently evolved “young” proteins compare to them, and whether random, young, and canonical sequences can interconvert along evolutionarily plausible paths, and which biophysical properties distinguish or link these sequences. Here, we analyse naturally occurring *de novo* proteins from yeast and flies, which originate from non-coding DNA and thus have experienced limited evolutionary selection. They serve as a model for examining the relationships between young *de novo* and intergenic proteins, older canonical proteins, and their randomized counterparts. Because *de novo* and randomized sequences lack detectable homology, we use an alignment-free k-mer-based distance approach. Randomization shifts distance distributions toward expected random behaviour in all classes, but natural, non-randomized sequence classes remain distinct, indicating non-random residue organization. Each class exhibits characteristic k-mer patterns, with *de novo* proteins clearly separated from both canonical and all randomized sequences. Sequences bridging these classes are frequently predicted to contain transmembrane helices. *De novo* proteins are thus not random samples of sequence space. Instead, they occupy constrained yet evolutionarily accessible regions defined by residue order and biophysical constraints, suggesting a plausible pathway for the emergence and diversification of new proteins.

**Significance Statement:** Despite the vast combinatorial potential of amino acids, evolution has produced only a limited repertoire of canonical proteins with conserved structure and function. How evolutionarily young proteins relate to older canonical proteins, and whether the sequence space between them is traversable, remain unclear. Here, we decompose canonical proteins, intergenic sequences, and recently emerged yeast and fly *de novo* proteins, together with randomized controls, into short, interpretable fragments (k-mers) and compare them using alignment-free distances. *De novo* proteins are markedly distinct from both randomized and canonical sequences. Notwithstanding their evolutionary distance, sequences are connected by stepwise paths comprising bridge sequences, often enriched for low-complexity motifs and transmembrane helices, connecting disordered and structured regions of sequence space.

## Introduction

The concept of protein sequence space describes the set of all possible amino acid sequences that could, in principle, form a protein [1]. Numerically, this high-dimensional space is astronomical: a protein of 100 amino acids could form 20^100^ possible sequences. In practice, however, the sequence of a protein is constrained by foldability [2], solubility [3], avoidance of aggregation [4], structural stability [5, 6], catalytic activity [7], epistasis [8], evolutionary trajectories [9, 10], and the limited evolutionary time to explore sequence space [11]. As a result, the set of sequences which have been evolutionary realised and experimentally determined represents only a minute fraction of the theoretical sequence space [12, 13]. Most known, evolutionarily “old” proteins (hereafter referred to as canonical proteins) occur in sets of evolutionarily related, highly similar sequences, i.e. protein families, that occupy distinct evolutionarily constrained regions of sequence space [14–18]. What remains unclear is how these constrained regions relate to the vastly larger space of weakly constrained or unconstrained sequences - such as evolutionarily “young” proteins and random (amino acid–shuffled) sequences- and whether, and by what routes, entirely new proteins can emerge from this background. Whether such emergence is possible depends on how canonical proteins, random sequences, and evolutionarily young proteins differ from each other, and on how navigable sequence space is between them; characterising these differences and quantifying that navigability is therefore central to understanding the limits of protein novelty. The “near-random” perspective of canonical proteins is supported by findings that sequence patterns of hydrophobic and hydrophilic amino acids - which are essential for structure formation - are similar between canonical and random sequences [19–25] and that some random proteins are predicted to form tertiary structures [26–28]. Likewise, *in vitro* experiments have shown that random sequence libraries can yield small fractions of folded and functional proteins [29, 30]. Such random proteins can, for example, rescue metabolic defects [31], confer toxin resistance [32], anti-phage functions [33], and catalyse biochemical reactions [34].

Moreover, many library-derived random sequences are soluble and exhibit defined secondary structure, suggesting that the capacity for folding and solubility may be relatively common even among random sequences, at least within crowded intracellular environments [35–39].

In support of the view that canonical proteins are “non-random”, biophysical and evolutionary constraints were found to leave characteristic signatures that sharply distinguish them from unselected random sequences [2, 40, 41]. This perspective is consistent with earlier hypotheses that random sequences are unlikely to produce biologically active or stably folded proteins [42, 43]. Computational analyses of hydrophobicity, hydrogen bonding, and electrostatic interactions indicate pervasive selective pressures that shape protein stability and constrain sequence evolution, supporting the non-random view [40, 44]. Simple biophysical models and information-theoretic approaches likewise support that natural proteins occupy discrete regions separated from random sequences [15, 45, 46]. These findings underpin the enduring metaphor of sequence space as an ocean dotted with “islands” of canonical protein families with largely conserved sequences, separated widely by non-functional random sequences [14, 15, 17, 47–51]. Although many protein families share measurable ancestry, sequence similarity, or structural features, they may remain widely separated in sequence space [52–54]. However, the distances separating canonical proteins from random and evolutionarily young proteins in sequence space remain unknown.

*De novo* emerged proteins occupy a particularly informative position in sequence space between random sequences and canonical proteins. Emerging from previously non-coding DNA and having experienced only limited evolutionary refinement, *de novo* proteins [55] represent some of the youngest naturally occurring proteins and are considered a close biological counterparts to random sequences [27]. Nonetheless, *de novo* proteins are subject to cellular expression, folding, and selection constraints [26, 27, 56–58]. This duality makes *de novo* emerged proteins an ideal system to determine connections between canonical and random sequences in sequence space.

Most known proteins, canonical proteins in particular, arise primarily through duplication of existing genes and subsequent sequence divergence [59]. *De novo* genes, which encode *de novo* proteins, emerge when previously non-coding DNA becomes transcriptionally active and produces a translated open reading frame (ORF) [60]. Initially considered exceptional, *de novo* genes have now been identified across eukaryotes, with several involved in key biological processes such as spermatogenesis in flies [61] and cell-cycle regulation in yeast [62]. *De novo* emerged proteins often exhibit high intrinsic disorder, marginal stability, and aggregation propensities, reminiscent of random sequences [26, 57, 63]. Some cases, however, exhibit secondary structures [61, 62, 64, 65] and others were predicted to integrate into membranes [66, 67].

*De novo* emerged proteins are often short, rapidly evolving, and enriched in low-complexity and intrinsically disordered regions. Therefore, distance measures that depend on homology, such as alignment-based methods, are inappropriate for comparing *de novo*, canonical, and random proteins [68–70]. Alignment-free, k-mer-based approaches provide an efficient alternative because they compare sequences without requiring positional homology and can capture local sequence organization even when global sequence similarity is absent [71–73]. Here, we use the alignment-free, k-mer-based tool SHARK-Dive (Similarity/Homology Assessment by Relating K-mers) [74] on naturally occurring *de novo* proteins from yeast and flies together with intergenic ORFs, canonical proteins, and their randomized counterparts, which we generate by maintaining amino-acid-composition and length distributions. We first ask whether natural classes are closer to being “near-random” or “non-random” by quantifying differences in residue order between natural and randomized classes, independently of length and composition. Next, we examine where *de novo* proteins typically lie in sequence space and whether they represent evolutionary transition points along paths from non-transcribed, near-random genomic sequences to canonical proteins. Finally, we characterize the biophysical features of k-mers that distinguish or are shared between classes.

## Results

We compared naturally occurring *de novo* proteins (DN), intergenic ORFs (Int), and conserved canonical proteins (Con) from fly and yeast, alongside length- and amino-acid-composition matched randomized controls (X-R) (Methods; Fig. 1; SI Appendix Fig. S1; Table S1). To quantify pairwise distances without requiring homology, we used SHARK-Dive [74]. Each sequence is decomposed into k-mer substrings of length *K* = 1–10. Pairwise comparisons across all k-mers are aggregated into a single scalar distance (Fig. 1C; Eq. 1). In SHARK-Dive [74], at *K* = 1 and *K* ≥5, distances are weighted by a physicochemical substitution matrix [76] (SI Appendix Table S2), combined with adaptive thresholding that retains meaningful k-mer signals at *K* ≥ 4. SHARK-Dive [74] thus overcomes the loss of sensitivity which is typically associated with k-mer methods at higher *K* [73]. By decomposing sequences into k-mers, we conceptually embed each sequence in a high-dimensional feature space defined by its k-mer composition, with pairwise distances quantifying geometric separation in a compositional sequence space rather than minimal mutational steps or alignment-based evolutionary divergence. This approach Offers a clear advantage over machine learning approaches because characteristic features of k-mers can be analysed and interpreted directly in a biophysical framework [77].

**Figure 1.**
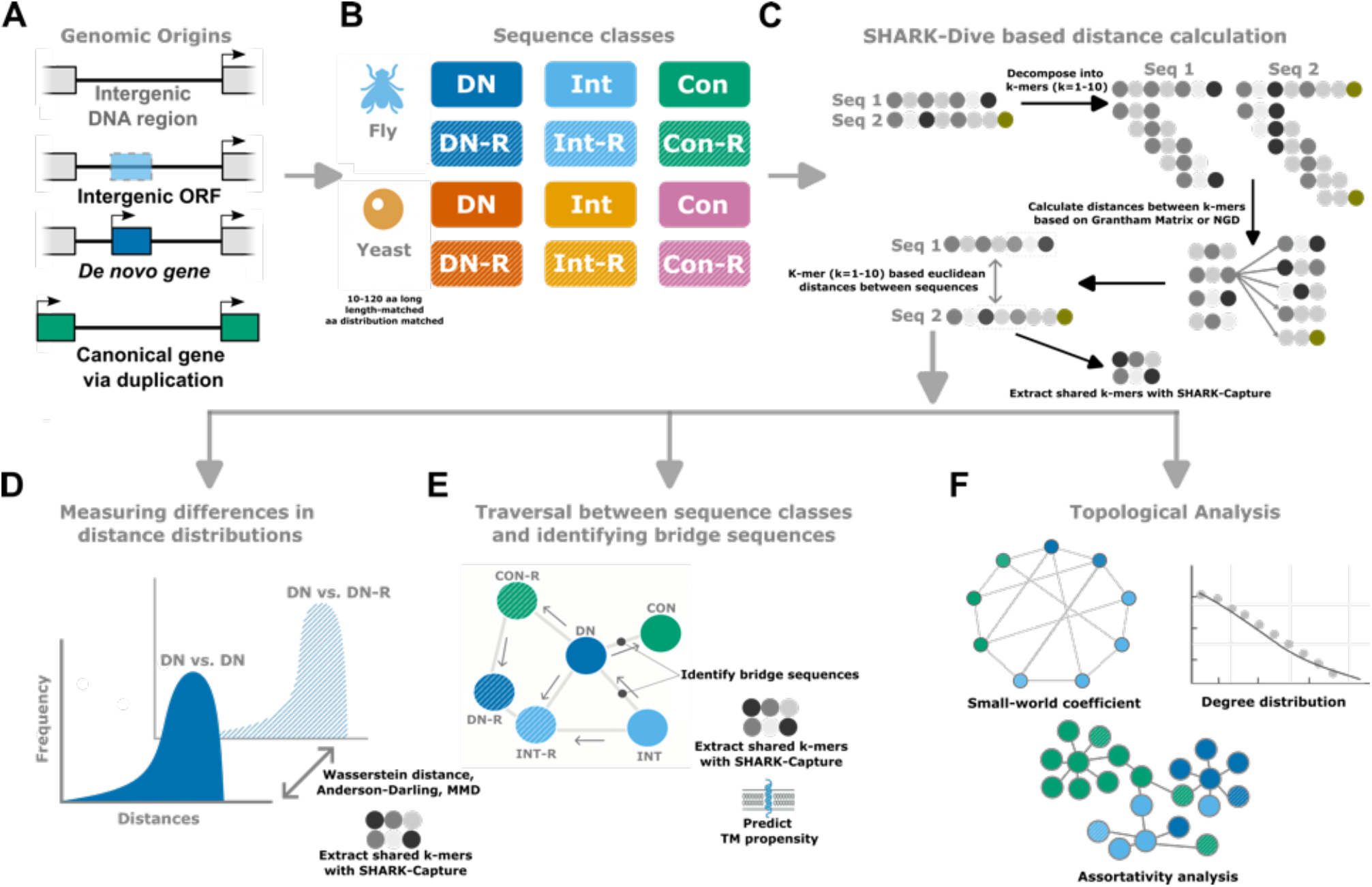
Workflow for measuring distances between young and old proteins using k-mer-based distances. (**A**) Genomic origins of datasets are intergenic (Int) ORFs (unannotated ORFs), *bona fide de novo* (DN) genes that emerged from formerly non-coding DNA and lastly canonical (Con) genes that emerged via duplication & divergence. (**B**) Sequence datasets comprise *de novo* (DN), intergenic (Int), and conserved (Con) proteins from fly and yeast, with length- and amino-acid-composition matched randomized controls (X-R). (**C**) Pairwise distances are computed by SHARK-Dive [74] derived k-mer distances (*k* = 1-10), combined into a unified distance for all-against-all comparisons (Panel adapted from Chow *et al.* 2024). (**D**) Downstream analyses include distributional comparisons of distances using Wasserstein distance, Anderson-Darling and MMD. (**E**) Identification of enriched k-mers using SHARK-Capture [75], traversal/bridge identification in a sparse k-NN graph, and (**F**) Global network topology (small-world structure, degree distributions, assortativity).

### Residue order separates all natural protein classes from randomized controls

We ask whether shared residue order brings natural sequences closer to one another than to randomized sequences in which that residue order is broken, i. e. does residue order create non-random local sequence patterns? To determine whether residue order contributes information beyond amino-acid composition, we compared for each natural class (DN, Int and Con) and organism (yeast and fly) the distribution of pairwise distances within natural sequences to the distribution of distances between each natural class and its randomized (-R) counterpart, using k-mer-based scalar distances (Eq. 1; SI Appendix Figs. S2 & S3) computed with SHARK-Dive [74]. If scrambling changes the distribution of pairwise distances, residue ordering contributes non-random local sequence patterns that are not explained by composition alone. The magnitude of this change therefore provides a measure of sequence-order information independently of amino-acid frequencies. To assess the magnitude of change in distance distributions, we used 1-Wasserstein (*W*_1_, Earth Mover’s distance) (Eq. 2), k-sample Anderson–Darling (AD) statistic (see Methods) and squared Maximum Mean Discrepancy (MMD^2^) (Eq. 3) to compare the distribution of distances with the corresponding distribution of distances when one of each pair was scrambled. The results are tabulated in Fig. 2. We repeated these calculations for three different sequence length bins and all possible combinations of pairs (10-50, 50-80, 80-120 aa; SI Appendix Table S3). *W*_1_ weights bulk and tail shifts equally and can therefore mask mass redistributed into the tails; because it also inherits the units of the SHARK distance, whose support widens at short sequence lengths, its magnitudes are not comparable across length bins. To distinguish a bulk shift from a tail-specific signature, and to obtain a length-comparable metric, we used the AD statistic (Fig. 2; see Methods), which emphasizes deviations in the distribution tails and has a known null distribution. Overall, and within each of the three length ranges, the W1 and AD tests indicated significantly larger deviations from randomized controls in fly DN and Con sequences than in Int sequences (Fig. 2; SI Appendix Table S3). Across length ranges, this trend was assessed using the AD statistic, which increased with sequence length in all three comparisons (SI Appendix Table S3). Interestingly, the differences from random in the yeast dataset were similar for all three sequence comparisons, increasing with sequence length for DN and Con but reaching a maximum at 50-80 aa for Int sequences (See SI Appendix: Table S3). Both the Wasserstein-1 and Anderson–Darling statistics quantify the magnitude of the change induced by scrambling rather than its direction. They therefore measure the extent to which residue ordering aKects the distribution of pairwise distances, irrespective of whether scrambling increases or decreases those distances. Because our objective is to quantify deviations from compositional randomness rather than determine whether scrambling increases or decreases pairwise distances, we interpret these statistics as measures of effect size rather than directional change. Neither *W*_1_ nor AD attaches a clean formal *p*-value to a single pair-of-pairs comparison, so we additionally asked whether the natural distance distribution is statistically distinguishable from its randomized counterpart at all. The MMD^2^ provides exactly this through a permutation test in which the binned counts of the two samples are reshuffled under the null hypothesis that they are drawn from the same parent distribution (Fig. 1; Methods; Eq. 3; SI Appendix Table S3). Of 72 comparisons (18 overall +54 length-binned), 69 reject the null at BH-adjusted *p <* 0.05; the three that do not reject the null are all comparisons of fly 10-50 aa Int sequences (DN-vs-Int to DN-vs-Int-R, Con-vs-Int to Con-vs-Int-R, Int-vs-Int to Int-vs-Int-R), with *W*_1_ shifts correspondingly small (↑ 0.006; SI Appendix Table S3). Biologically this means short, intergenic fly sequences carry no detectable residue-order structure: shuffling them does not move the resulting distance distribution, so short, intergenic fly sequences are “near-random” with respect to the SHARK metric, and any cross-class pairing anchored on them inherits that randomness. Taken together, randomization of residue order shifts the k-mer distance distribution of essentially every sequence class in both species - most strongly for longer fly DN/Con sequences and for yeast Con sequences, in which the shift grows with sequence length in both species.

**Figure 2.**
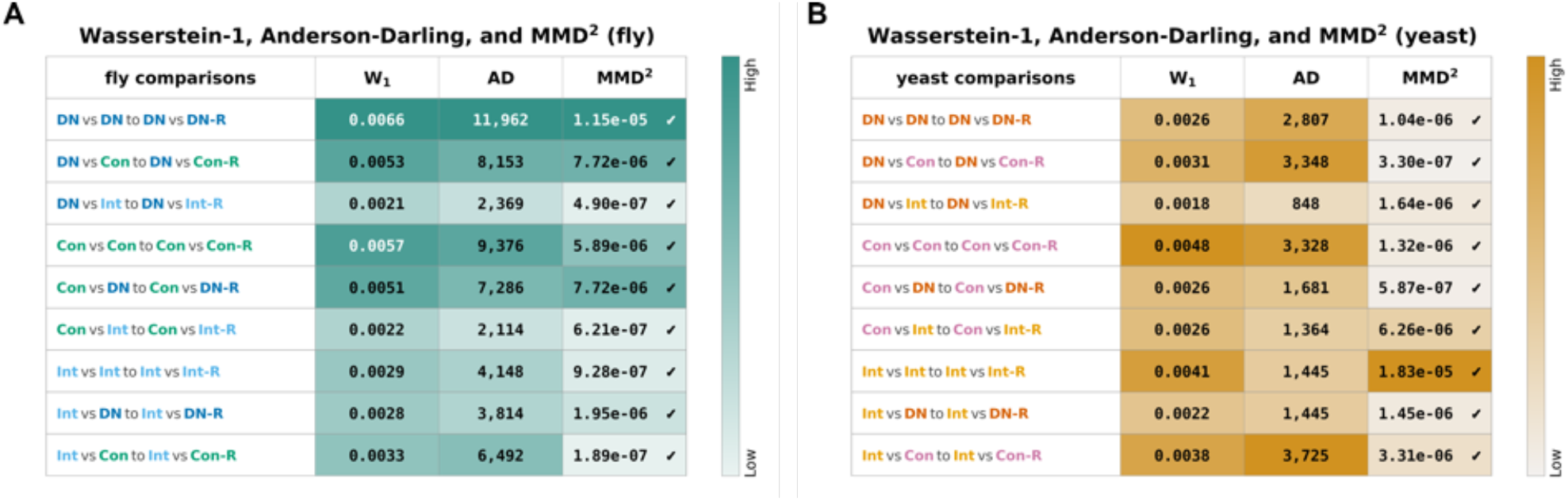
Randomization reshapes pairwise distance distributions in fly and yeast. Wasserstein-1 distance (*W*_1_; Methods; Eq. 2), k-sample Anderson-Darling statistic (AD; Methods), and squared Maximum Mean Discrepancy (MMD^2^; Methods; Eq. 3) between pairwise distance distributions of each natural-class comparison and its randomized control (X-R) in fly (**A**) and yeast (**B**). For an ordered class pair (A*, B*), the three metrics quantify how the natural distance distribution *P_A,B_* differs from *P_A,BR_* , where *B* is replaced by its randomized counterpart *B_R_*; this includes within-class comparisons (e.g., *W*_1_(*P*_DN,DN_*, P*_DN,DN-R_)). *W*_1_ carries the units of the underlying SHARK distance; AD is a unitless standardised k-sample statistic; MMD^2^ values are shown together with where the null of identical distributions is rejected by the permutation test at Benjamini-Hochberg-adjusted *P <* 0.05 and × otherwise. Cells are coloured per metric, with each metric independently min-max normalised. SI Appendix Table S3 lists the same three metrics across overall and three sequence-length bins (10-50 aa, 50-80 aa, 80-120 aa); SI Appendix Figs. S2-S3 show raw distance distributions between all classes.

### Enriched k-mers reveal constrained local sequence structure beyond composition

We next aimed to identify the sequence motifs underlying these non-random residue-order patterns that are lost upon randomization. Analogous to our k-mer–based scalar distances, we used SHARK-Capture [75] to extract the top ten enriched k-mers (*K* = 3-10) for every class (Methods). In fly, DN and Con are enriched for low-complexity, repetitive motifs dominated by glutamine (PolyQ; e.g., QQQ–QQQQQ) and leucine (PolyL; e.g., LLLL), as well as mixed motifs. Specifically, DN shows enrichment of LLP and LPL, whereas Con is enriched for QLL and KLL (Fig. 3). PolyQ motifs are absent from Int, although PolyL (LLLL) is also enriched in Int. The depletion of these motifs upon randomization indicates that their enrichment reflects residue-order–dependent sequence features rather than amino-acid composition alone (Fig. 3; SI Appendix Figs. S1 & S4). In contrast, several hydrophobic motifs (e.g., LLA, LLL, LLS) appear in both natural and randomized sets, consistent with being largely composition-driven rather than class-specific. Notably, Con-R contains hydrophobic k-mers such as LLR, LRL, and RLL, indicating that local hydrophobic motifs can also arise by chance under randomization. Yeast showed a less pronounced enrichment: DN and DN-R repeatedly contained polar S/T-rich motifs (Fig. 3; SI Appendix Fig. S4). By contrast, Con and Con-R contained more S/L-rich motifs. In yeast, residue patterns contribute less to sequence differentiation between classes, while amino-acid composition alone can give rise to hydrophobic motifs, such as LLS, LLK or ILL. Together, k-mer enrichment supports a simple distinction: hydrophobic fragments largely reflect background composition and can be enriched also in randomized sequences, whereas low-complexity motifs - most clearly PolyQ and PolyL in fly - are lost under randomization and are enriched in natural classes.

**Figure 3.**
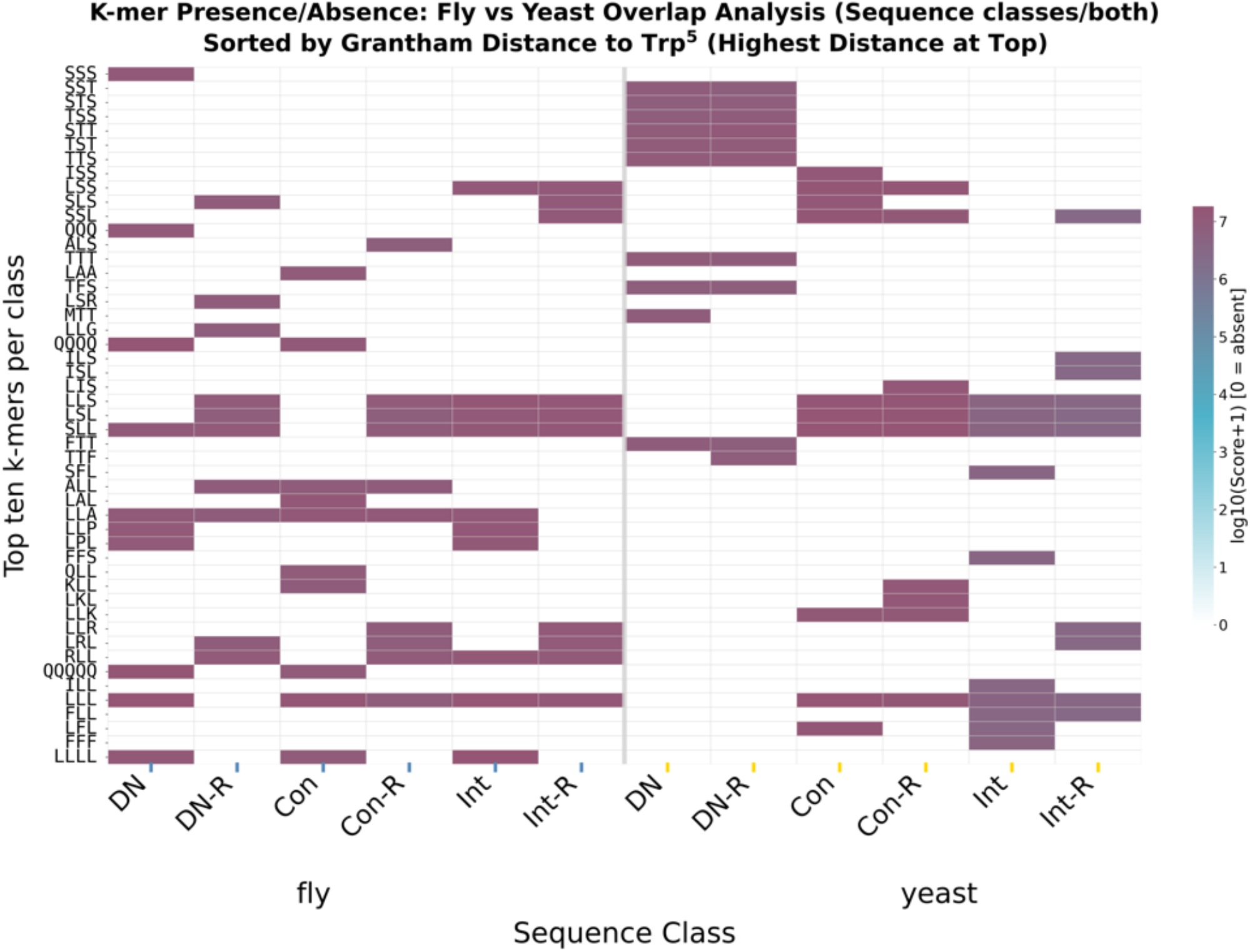
Enriched k-mers across sequence classes and species reveal distinct motifs that can be disrupted by randomization but some hydrophobic motifs do appear enriched in randomized sequences. Presence/absence heatmaps of top ten k-mers (*K* = 3-10) across natural classes and matched randomized controls in fly (left) and yeast (right). Ordered by Grantham distance [76] to Trp^5^. SI Appendix Fig. S4 for length binned analysis of motifs.

### Natural classes are easy to reach, randomized classes are easy to leave

We next asked whether sequence class differences relate to distances and connectivity of their sequences in sequence space. We used traversals starting from every sequence on a nearest neighbor graph (Methods; Eq. 4) to characterize each class by its accessibility, i.e., how easily its sequences can be accessed from other classes, and traversability, i.e., how readily sequences can traverse to other classes (Fig. 4; Methods; Eq. 6). Across yeast and fly, all randomized classes (X-R) were consistently highly traversable but weakly accessible (Fig. 4; SI Appendix Figs. S5 & S6), indicating that randomized sequences tend to locate in regions of sequence space that are easy to “exit” but are rarely accessed when traversing away from natural classes. In fly, natural classes were clearly separated, with Con showing high accessibility and low traversability, consistent with dense local neighborhoods of canonical proteins. On the other hand, DN occupied an intermediate region relative to Con and Int (Fig. 4A). In yeast, class separation was weaker and DN lay closer to Int than to Con, consistent with a more compositionally similar landscape between natural classes (Fig. 4B). These patterns show that non-random residue patterns and class-specific sequence constraints shape not only pairwise distances but also the navigability in sequence space. In both taxa, DN is substantially more accessible than X-R, positioning DN between Int and Con, while Int is the natural class closest to X-R.

**Figure 4.**
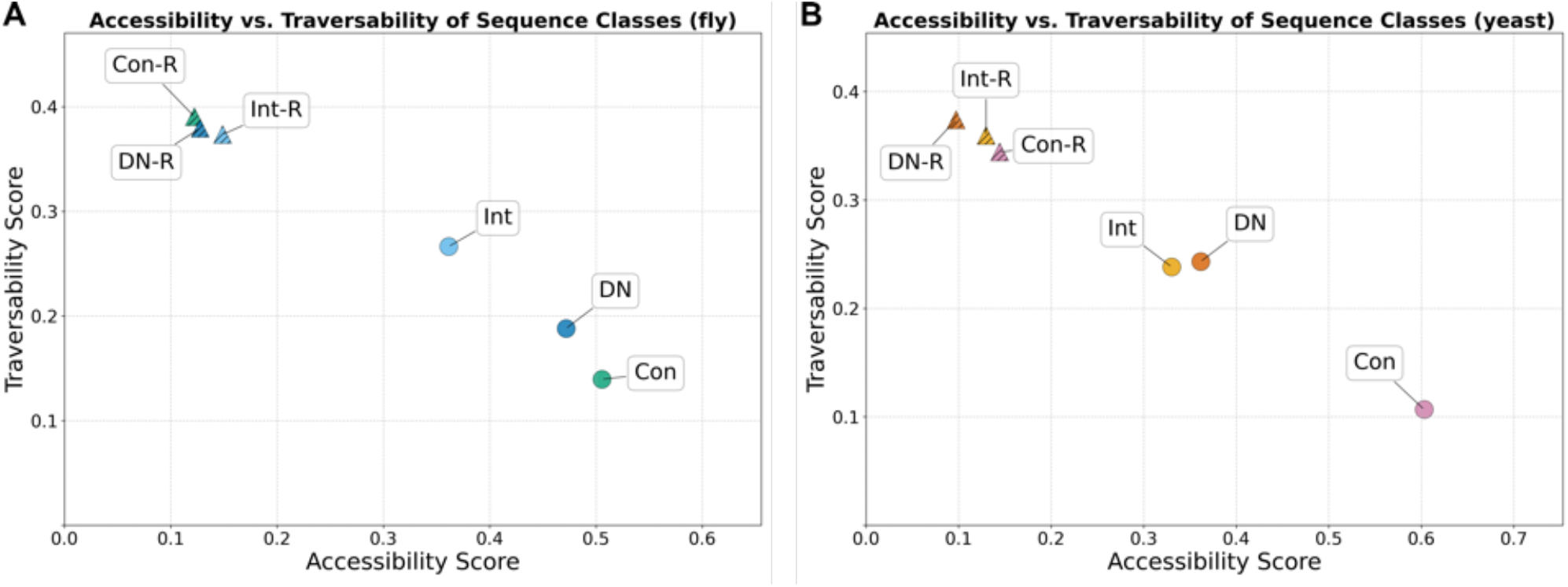
Class-level accessibility and traversability in sequence space reveals that regions of sequence space occupied by randomized sequences are hard to reach but easy to leave. Accessibility (x-axis) and traversability (y-axis) (Methods; Eq. 6) scores for each sequence class in (**A**) fly and (**B**) yeast, computed from greedy traversal success across ordered class pairs in a sparse k-NN graph (Methods; Eq. 4). Points show natural classes (DN, Con, Int) and their matched randomized controls (X-R) scaled by bridge score.

### Bridge sequences are rare and enriched for transmembrane features

Building on the nearest-neighbor traversal, we next examined the local arrangement of classes and their sequences in sequence space. Sequence classes could be intermingled or instead form compact, contiguous regions, even though their constituent protein families likely have largely independent evolutionary origins. If classes do occupy dense regions, then “bridge sequences” should exist that permit stepwise nearest-neighbor moves through sequence space and connect members of different classes [78, 79]. We therefore calculated a bridge score (Methods; Eq. 8), describing how often sequences appeared as intermediate nodes on traversals and analysed the predicted structural properties of the top ten of these bridge sequences per taxon (Methods; Fig. 5; SI Appendix Fig. S7-S10). The top ten identified bridge sequences include representatives from all natural classes in both taxa, and only two of them are randomized sequences. Bridge sequences were strongly enriched for predicted transmembrane (TM) regions (Fig. 5). In fly, 8 of 10 bridge sequences contain at least one TM helix. In yeast, 7 of 10 bridge sequences contain → 1 TM-helices. Consistent with this, bridge-connected sequences were enriched for hydrophobic k-mers (e.g., I/L/V-rich fragments) that typically promote membrane localization [80] and hydrophilic serine repeats, which have been shown to be largely neutral and to reduce aggregation [81] (SI Appendix Figs. S7-S10; Tables S4-S6). AlphaFold3 frequently predicts α-helices in those bridge sequences, and many contain low-complexity repeats (Fig. 5; SI Appendix Fig. S7), linking bridge sequences to concrete biophysical sequence features. This is further corroborated in all classes, except in DN-R, by overall positive correlation between TM-propensity and bridge score (Methods; Eq. 8), with the strongest association in fly Int (Spearman ρ = 0.180, *p <* 10^−15^) and yeast DN (ρ = 0.153, *p <* 10^−9^) (SI Appendix Table S6). In contrast, DN-R showed no association between TM-propensity and bridging in either species, suggesting that TM-promoting features (helices, motifs) have been shaped by selection in DN, which are disrupted by randomization in DN-R. Taken together, these results show connectivity between Int, DN, Con and randomized counterparts is mediated by rare bridge sequences which are enriched for specific, biophysically interpretable features such as low-complexity sequence repeats and hydrophobic, helix-forming segments with elevated TM-propensity.

**Figure 5.**
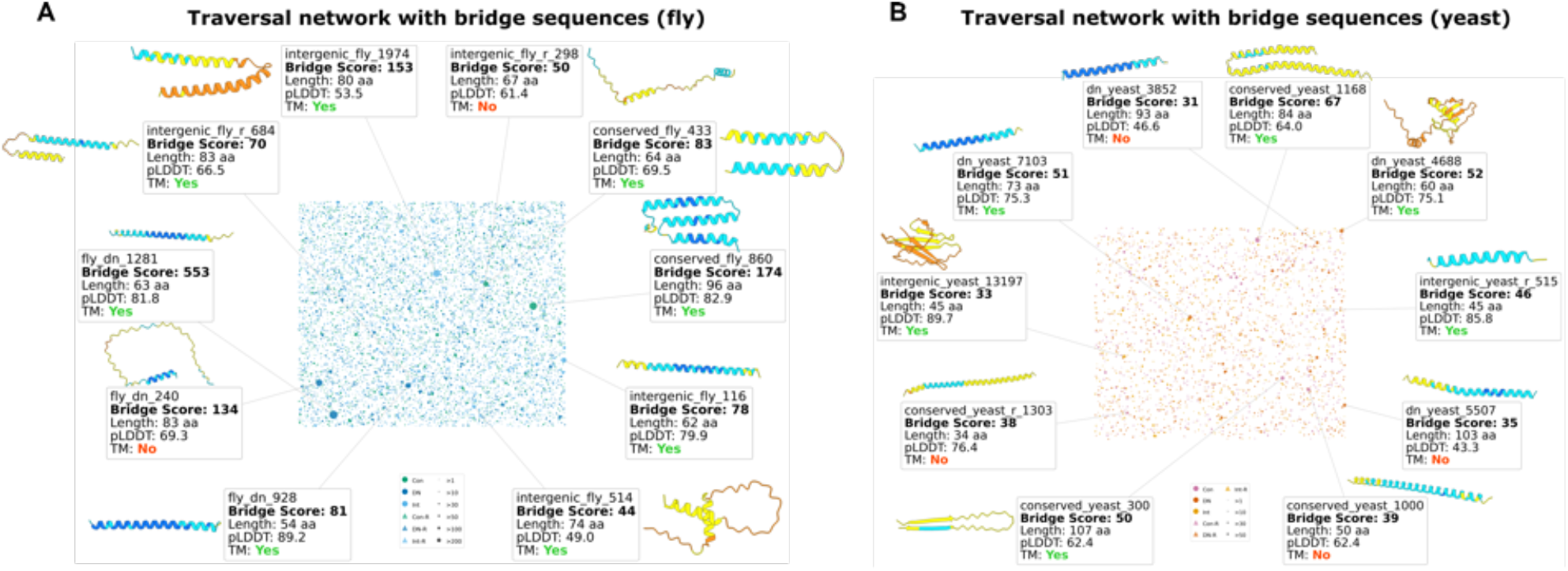
Traversal bridge networks and AlphaFold3 structural predictions of top bridge sequences reveal sequences enriched for transmembrane helices. The bridge network was visualised using the Fruchterman-Reingold spring layout [82] applied to the greedy nearest-neighbor traversal paths that cross at least one bridge node. Nodes represent sequences with a significant bridge score (observed nearest-neighbor traversal count → 1) or connected sequences. (**A**) for fly, (**B**) for yeast. The ten sequences with the highest bridge scores are annotated with labels indicating bridge score, sequence length (aa), predicted local distance difference test score (pLDDT), and transmembrane prediction (TM). For each of the ten top bridge sequences, the adjacent panel shows the AlphaFold3-predicted tertiary structure, coloured by per-residue pLDDT confidence score. SI Appendix Figs. S5 & S6 show transition probabilities and path lengths. SI Appendix Figs. S7-S10 show Bridge scores, identified motifs, sequences, TM-proportions, and sequences connected by bridge sequences. SI Appendix Table S4-S6 show statistical testing of bridge sequences and statistical correlation between bridge score and TM prediction.

### Global topology explains clusters connected by rare hubs

To characterize the large-scale organization of sequence space, we constructed k-nearest neighbor networks (k = 3, 5, 10) from pairwise sequence distances and analysed their topology both overall and within each class (Methods). Both fly and yeast networks exhibited, overall and in the natural classes, large small-world coefficients, indicating simultaneously high clustering and short average path lengths relative to random networks. In contrast, small-world coefficients were low in X-R classes, indicative of only sparse clustering and longer path lengths (Fig. 6A; SI Appendix Fig. S11). Degree assortativity - the tendency of nodes to connect to others of similar degree - was near zero to slightly negative, indicating that high-degree nodes (sequences with many close neighbors) preferentially connect to low-degree nodes (sequences with few close neighbors) rather than to each other (Fig. 6C). Degree distributions were heavy-tailed, confirming that a small subset of sequences acts as highly connected hubs within a majority of sparsely connected nodes (Fig. 6B, D). Class-level patterns were consistent with the traversal results: natural classes, and Con in particular, form clusters, whereas networks of randomized classes showed substantially reduced clustering and longer average path lengths closer to random expectation. Together, these topology measures show that sequence space combines local density with global reachability: class-specific clusters are linked by a small number of highly connected intermediates, consistent with the bridge sequences identified above.

**Figure 6.**
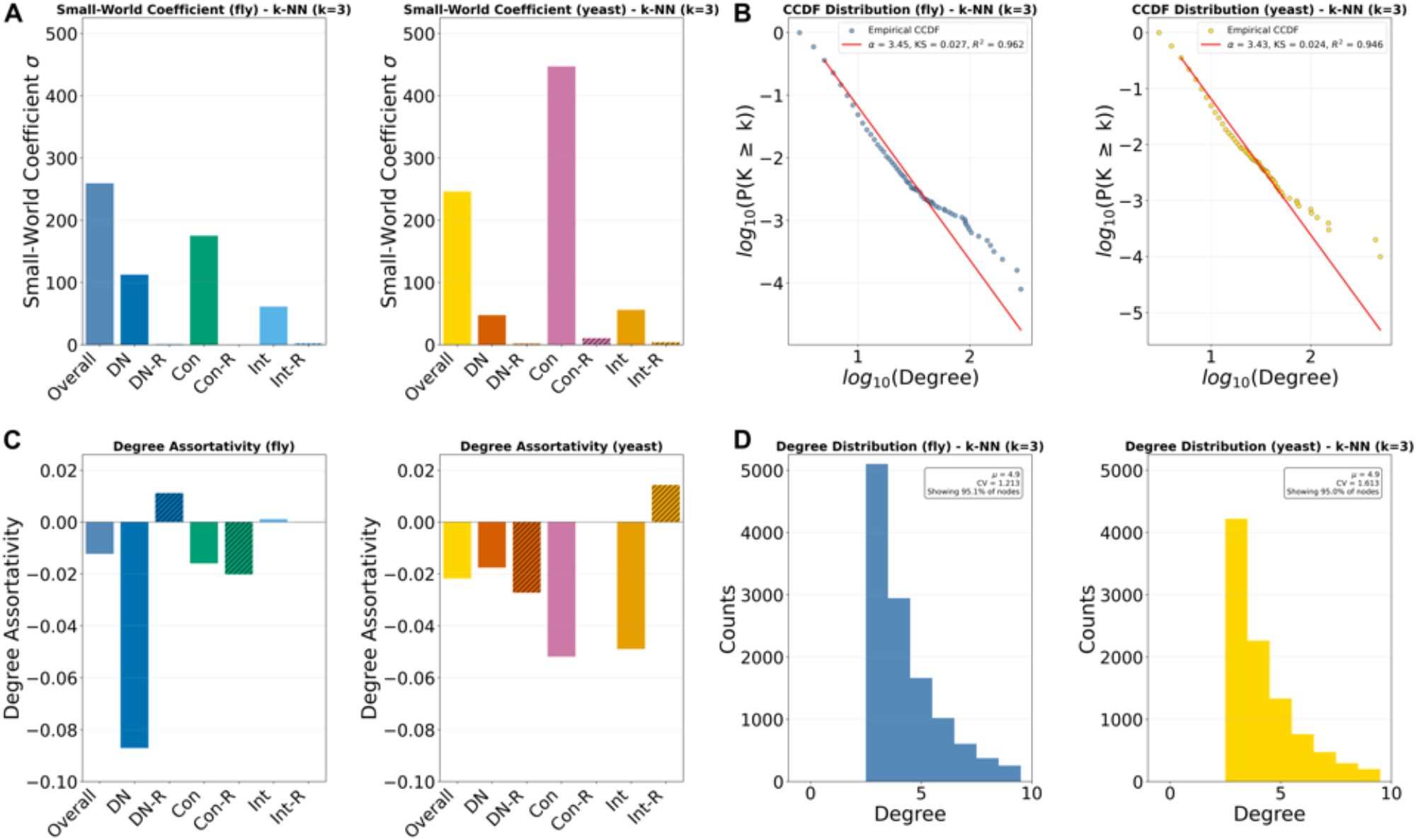
Global topology of k-NN sequence-space networks (k=3) reveals dense clusters that are connected by hubs. (**A**) Small-world coefficient (σ) for each class-specific k-NN network in fly (left) and yeast (right). (**B**) Complementary cumulative degree distributions (CCDFs) for the overall fly and yeast networks shown on log-log axes with discrete power-law fits (fit parameters reported in the panels). (**C**) Degree assortativity (*r*) for the same networks. (**D**) Corresponding degree histograms for the overall fly and yeast networks (k=3), illustrating degree heterogeneity. SI Appendix Fig. S11 for analysis of k-NN= 5, 10.

## Discussion

### Randomization reveals a structured and navigable protein sequence space

Our results consistently show that protein sequence space is non-random and non-uniform beyond amino-acid composition alone. We demonstrate, using *W*_1_ combined with MMD and AD, that residue order contributes substantially to how natural classes are positioned in sequence space: shuffling residues alters distance distributions in all sequence classes, most strongly in longer fly DN & Con and in yeast Con - signals that grow with sequence length in both species - and most weakly in Int, where short fly intergenic sequences are statistically indistinguishable from random (Fig. 2; SI Appendix Table S3). In yeast, intergenic sequences appear close to randomized controls in bulk *W*_1_ comparisons but retain localised ordering-related features - revealed by MMD - that distinguish them from coding sequences and disappear upon randomization. One explanation is that many yeast intergenic ORFs are derived from genomic regions with coding-like composition (e.g., untranslated regions, transposon fragments, or cryptic ORFs) [66, 83, 84] (SI Appendix Fig. S1; Table S1). Randomization removes these weak signals and shifts Int toward the randomized controls. Traversal, bridge, and network analyses reinforce this picture. Con is highly accessible and strongly clustered, Int is closest to randomized classes, and DN falls consistently between them (Fig. 4 & Fig. 6). This intermediate position is further supported by a length dependency: short DN sequences resemble Int and randomized sequences (low AD), while longer DN sequences diverge from random expectation (high AD; SI Appendix Table S3), consistent with longer *de novo* proteins accumulating more position-specific constraints over time [85].

### Motifs are lineage-specific: poly-Q in flies and composition-driven patterns in yeast

Motif enrichment separates two sources of signal: hydrophobic fragments that largely track background composition and persist after randomization, and low-complexity motifs that are disrupted by shuffling and enriched only in natural sequences (Fig. 3). In fly, the clearest example is poly-Q, which is common in metazoan regulatory proteins and strongly linked to neuronal function and pathology [86], explaining its prevalence in fly but not yeast. The enriched poly-L runs (e.g., LLLL) promote α-helix formation and facilitate membrane integration [87], consistent with widespread helical structure across classes [27, 88] and the elevated membrane propensity of bridge sequences (Fig. 5). In yeast, many k-mers are enriched in the amino acids S, T and L, across natural and randomized classes (Fig. 3), indicative of yeast sequences being strongly composition-biased (SI Appendix Fig. S1; Table S1). Therefore, expected frequencies alone explain much of the observed k-mer signal (SI Appendix Fig. S1). Since hydrophobic motifs occur even in Int and X-R, short structure-forming motifs can arise by chance given the underlying amino-acid composition. This is consistent with experimental studies showing that random sequence libraries can yield folded proteins [29, 35, 38]. Our results also highlight an important role of repetitive, low-complexity motifs in shaping natural sequence space. Because these motifs depend on residue order, they are disproportionately disrupted by randomization: shuffling breaks repetitive patterns and consequently removes intrinsically disordered segments while preserving overall composition. This Offers a simple explanation for why randomized sequences are predicted to be less disordered than their natural counterparts (SI Appendix Fig. S12) [26, 57, 63, 89]: randomization disrupts not only structure-forming fragments but also low-complexity regions that promote disorder, shifting sequences toward more mixed, less-repetitive local patterns [38].

### Membrane association is a robust route for bridging sequence space

Bridge sequences connecting different classes in sequence space are strongly enriched for transmembrane helices in both species (Fig. 5). This enrichment likely reflects the capacity of membrane association to stabilise otherwise marginally stable peptides comprising amphipathic or TM-helices, increasing both their physiological and evolutionary persistence. This aligns with predictions that *de novo* proteins have high propensity to interact with membranes and condensates [27, 66, 67], and with the role of the identified L-rich motifs in membrane integration [80]. Membranes act as scaffolds that promote helical structure in otherwise flexible proteins and can buffer aggregation risk by confining proteins into specific cellular contexts [80]. A TM-segment is a short, compositionally simple feature that can arise from many sequence variants, explaining why such segments appear across all sequence classes, including Int, DN, and X-R. Critically, the positive correlation between TM-propensity and bridge score in all classes (except DN-R) indicates that TM-promoting features have been shaped by selection and are disrupted by randomization (SI Appendix Table S6). Our results therefore suggest that the path from non-coding sequences to retained proteins may pass through bridge sequences characterized by motifs facilitating membrane association. This matches the biophysical view of protein evolution as “locally walkable” but “globally constrained” by stability, misfolding, and aggregation [13, 90]: evolution can only move through regions where intermediate sequences remain viable. A comparable link between membrane association and evolutionary intermediacy has been proposed for the prion protein, which may retain an evolutionary “imprint” of TM-ancestry and persist in conformational exchange between soluble, disordered and membrane-associated, structured states [91]. Given the high disorder levels and enriched low-complexity motifs in natural sequences, the helical, TM-containing bridge sequences may thus connect disordered with structured regions of protein sequence space.

### Limitations and scope

Our analyses describe protein sequence space using alignment-free k-mer distances, which capture short-range organization but do not explicitly model long-range tertiary interactions or functional activity. Traversal analyses rely on greedy nearest-neighbor walks and therefore reflect local navigability rather than optimal evolutionary paths. Although we refer to these traversal metrics as accessibility and traversability following previous graph-theoretic terminology, they quantify properties of the inferred sequence-space network rather than evolutionary accessibility directly. Differences may also reflect local sequence density or clustering under the chosen network construction. In addition, our analyses are limited to sequences between 10 and 120 amino acids. This range was chosen to avoid becoming dominated by the combinatorial vastness of sequence space while still covering the length scale of motifs and domains. Our conclusions are also based only on two taxa, which are however evolutionarily very distant from each other. Despite these limitations, the convergence of distance-based, traversal, motif, and topological analyses suggests that the major patterns we identify reflect robust features of protein sequence space rather than artifacts. The non-random signals in Int and DN are consistent with two non-exclusive mechanisms - weak protein-level selection via noise-induced transcription [60, 92] and DNA-level constraints (nucleosome positioning, GC content, replication stability) that bias amino-acid composition independent of protein sequence [93]. These two mechanisms cannot be fully disentangled by sequence data alone. Lastly, we can only analyse a current snapshot of protein sequences and dense connectivity might be the result of limited time for proteins to diverge [11].

### Conclusions and outlook

Across two evolutionarily distant taxa, our results show that protein sequence space is structured beyond amino-acid composition alone. By using an alignment-free, k-mer framework, we were able to quantify distances in sequence space independently of detectable homology and extract underlying motifs. This allows us to assess non-random organization even among sequences that cannot be aligned and would therefore remain invisible to traditional homology-based approaches. We show that DN proteins are not simply natural random sequences. Instead, they occupy a constrained part of sequence space, between Con and Int, shaped by short-range sequence organization and often connected by a small set of membrane-associated intermediate bridge sequences. Their enrichment for TM-helices suggests that membrane association provides a recurrent route by which newly emerging sequences can persist and connect regions of sequence space. While natural sequences are non-random, some randomized sequences can still acquire properties similar to natural proteins, simply by chance drawn from a natural amino acid distribution. Our results provide a more nuanced view, beyond the proposed dichotomy of natural sequences being near-random or non-random. Natural sequences, including intergenic ORFs with the exception of very short ones (10-50 aa), show clear non-random patterns beyond amino-acid composition. At the same time, consistent with the near-random view, randomized sequences can still contain short motifs and associated properties: they can form structures, including predicted TM-helices, and in rare cases act as bridge sequences in traversal. This is consistent with the idea that many protein-defining features arise from basic physicochemical constraints of amino acids, and that evolution primarily finds, selects and stabilizes such properties rather than explicitly “building” them from scratch [19]. In this view, randomized sequences can occasionally resemble natural proteins because short motifs can emerge by chance given the composition, and these motifs can provide starting points for helix formation, weak interaction interfaces, or membrane integration. Our motif results support exactly this: much of the signal that separates classes is carried by small sequence features. Together, this suggests that the effective “building blocks” shaping sequence space are short, local patterns. Mapping and quantifying these patterns provides a practical way to compare natural and randomized sequences without homology, and Offers a path toward designing and testing sequences by combining motifs that move sequences into regions of sequence space beyond the canonical [10, 94, 95]. This perspective also places *de novo* emergence in a broader evolutionary context. Evolution works opportunistically with available material under constraint and history. *De novo* proteins, intergenic ORFs, and experimental random peptide libraries can all be viewed as sequence material on which selection can act. In this view, *de novo* and randomized sequences are not simply distant outliers in sequence space, but accessible starting points from which novel proteins may be evolved.

## Materials and Methods

### Sequence selection and generation

*De novo* (DN) proteins were taken for *Drosophila* from Heames *et al.* (2020) [96] and Peng & Zhao (2024) [88], and for yeast from Blevins *et al.* (2021) [97] and Wacholder *et al.* (2023) [92]. All sequences were filtered to 10-120 aa, canonical amino acids, and made non-redundant. Conserved proteins (Con) were sampled from reference proteomes to match the relevant length distributions (fly: combined proteomes of 12 *Drosophila* species; yeast: *S. cerevisiae* proteome), and DN entries overlapping these proteomes were removed. Intergenic ORFs (Int) for fly and yeast were obtained from Iyengar *et al.* (2024) [98], de-redundified, and sampled to match the input length distribution (fly: DN; yeast: Con). For each set we generated length- and amino-acid-composition matched randomized controls (X-R). A total of 12,414 fly sequences and 10,040 yeast sequences were analyzed. Dataset sizes and compositional properties are provided in SI Appendix Fig. S1; Table S1.

### Pairwise distance calculations between sequences

As alignment-free k-mer distances for *K* = 1-10, we used SHARK scores [74]. Briefly, SHARK-Dive calculates scores using a Grantham-weighted physicochemical distance [76] for *K* = 1 and *K* = 5-10 (SI Appendix Table S2), whereas for *K* = 2-4, it uses the Normalized Google Distance [99] and outputs a final classifier score. We re-inverted the SHARK scores to distances and combined them into one distance without the classifier score. Let *d_k_* (*i, j*) denote the distance between sequences *i* and *j* for k-mer size *K*. We then combined distances across all k-mer sizes using the Euclidean norm:

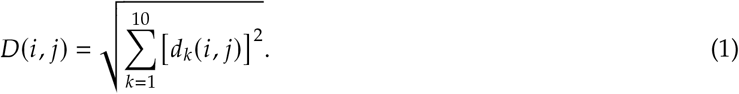

Shared k-mers (*k* = 3-10) were extracted with SHARK-Capture [75] (top ten consensus k-mers).

### Measuring distance diHerences between and within sequence classes

Pairwise-distance distributions were compared using the 1-Wasserstein distance (Earth-Mover distance) [100]:

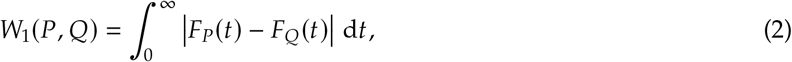

where the empirical cumulative distribution function (ECDF) *F_p_*(*t*) is the fraction of raw distances in sample P with value ≤ *t*; *W*_1_ is therefore the area between the two step-function ECDFs of the unbinned, raw samples. Two supplementary metrics provide orthogonal views: the k-sample Anderson-Darling test [101], an ECDF goodness-of-fit statistic that upweights deviations in the distribution tails where *W*_1_ averages them out, and the squared Maximum Mean Discrepancy (MMD^2^) [102], a kernel-based discrepancy whose permutation null yields a *p*-value that neither *W*_1_ nor AD can provide cleanly. MMD^2^ measures the squared distance between two distributions after mapping each into a feature space via a positive-definite kernel and comparing their mean embeddings; with a Gaussian kernel discretised on a shared 1-D support, this reduces to the closed form

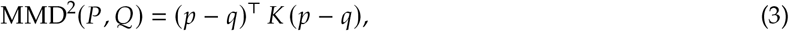

where *p*, *q* are bin-probability vectors on a 100-bin support shared by all comparisons within a given (species, length-bucket) and *K_ij_* = exp(–(*c_i_* – *c_j_*) ^2^ / (2σ^2^)is the Gaussian kernel at the bin centres *c_i_* , with σ fixed once per bucket as the median pairwise gap of those centres. Sharing the support and σ across pairs in the same bucket makes MMD^2^ values directly comparable. A permutation-test *p*-value is computed from 500 multivariate-hypergeometric resamples of the bin counts under H_0_ that the two samples share a distribution, taking 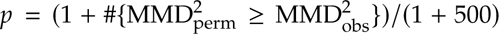. *p*-values are adjusted using the Benjamini-Hochberg false-discovery-rate (FDR) procedure [103].

### Traversal between sequence classes

For all sequences (*s*_0_), we performed a greedy nearest-neighbor walk *p* = (*s*_0_*,. . . , s_h_* ), to reach all other sequence classes by stepping to the single closest neighbor:

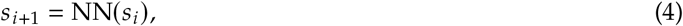

and terminating upon revisiting a node, reaching the target class, or exceeding 50 hops. For each successful traversal *i* to target class *T* we recorded number of hops *h_i_* and cumulative distance 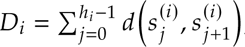, and report 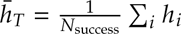 and 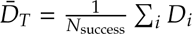. Across all attempts, traversed paths covered 6,764 of 12,414 fly sequences (54.5%) and 5,507 of 10,040 yeast sequences (54.9%) as non-start (visited) nodes; i.e. just over half of the sequence space was reached by at least one traversal, while every sequence served as a starting point. The set of sequences visited by successful walks was identical to that visited by all walks, indicating that failed traversals pass through the same hubs rather than exploring otherwise-unreached regions.

#### Attempt- and hop-normalised class reachability

For each class pair (*A B*), with 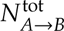 traversal attempts (including failures), we defined a hop-normalised success score:

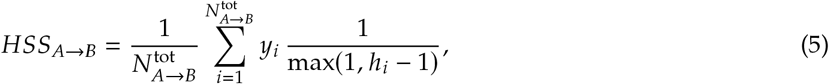

Where y_i_ = 1 if attempt *i* reached *B* (TargetReached=true) and 0 otherwise (so failed attempts contribute 0). Accessibility and traversability were then computed (excluding self-pairs) as:

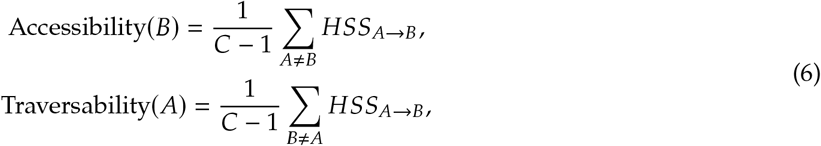

with *C* the number of classes. Although we record the cumulative path distance D_i_ , the reachability and bridge scores (Eqs.5-8) are normalised by hop count rather than by *̄D_i_* . Because each step moves to the single nearest neighbor, the per-step distances are narrowly distributed, so *D_i_* ≈ *̄d, h_i_* is to a good approximation a deterministic multiple of the number of hops and carries essentially the same information. Critically, per-hop distance does not separate successful from unsuccessful attempts, whereas hop count does; cumulative distance therefore contributes no independent signal about reachability. Hop-based normalisation is also unitless and independent of the absolute scale of the distance metric, making scores comparable across class pairs and robust to a few atypically long steps, which a distance-weighted score would over-emphasise. We thus report *D_T_* descriptively (SI Appendix Fig. S6) but base all scores on hops.

#### Bridge sequences and enrichment

Bridge candidates were ranked by raw intermediate-use counts across successful paths:

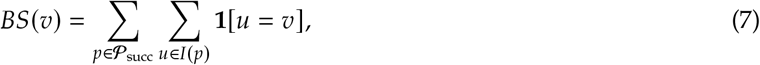

where *I*(*p*) are intermediate nodes (excluding start/target). For statistical testing we used a hop-normalised weighted bridge score:

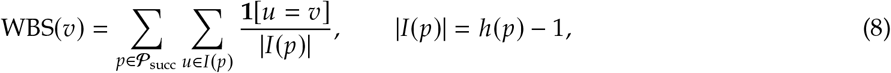

so each successful path contributes unit weight distributed across intermediates. Empirical nulls (*B* = 200) were generated by (i) degree-preserving edge rewiring of the 3-NN graph [104] and (ii) distance-based randomization sampling intermediates with probability ∝ 1/(*d*_start_ + *d*_target_) from the local 1-2 hop neighborhood. For each null, traversals were recomputed and WBS recorded. One-sided empirical *p*-values were:

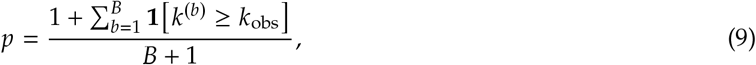

class-pair-specific *p*-values were combined per node using Fisher’s method, and FDR was controlled with Benjamini-Hochberg (*q ≤* 0.05).

### Structural predictions

Structure predictions were performed with AlphaFold3 [105] and visualized with ChimeraX-1.10 [106]. Disorder was predicted with PUNCH2-light [107] and compared across groups using Kruskal-Wallis tests followed by Dunn’s post-hoc tests with Holm correction. Transmembrane (TM) propensity was predicted with Phobius [108]. TM-enrichment in bridge sequences versus species-wide backgrounds was tested with one-tailed Fisher’s exact tests and one-tailed binomial tests with Benjamini-Hochberg FDR correction; 95% Wilson confidence intervals were used for TM-positive fractions. Associations between bridge usage and TM-propensity were assessed using Spearman rank correlations and Mann-Whitney U tests (TM-positive: → 1 helix), with Benjamini-Hochberg correction across tests (*q <* 0.05).

### Network construction for topology analysis

To avoid complete-graph artifacts from all-against-all distances, we constructed k-nearest-neighbor graphs by connecting each sequence to its *K* closest neighbors under *D*(*i,k*) (main analysis *k* = 3; robustness *k* = 5, 10, SI Appendix Fig. S11). Class-specific networks were analysed as induced subgraphs retaining only nodes from a given class and edges between them.

### Degree distribution analysis

Degree distributions were computed for each k-NN network. For node *i* with degree *k_i_*, we analysed the complementary cumulative distribution function (CCDF) and fit discrete power laws to the tail following Clauset *et al.* (2009) [109]. The lower cutOff *K*_min_ was chosen by minimizing the Kolmogorov-Smirnov (KS) distance; the exponent γ was estimated by maximum likelihood for *k_i_ ≥ K*_min_, with tail behaviour *P* (K ≥ *k*) ∝ *k^−^*^(*γ*–1)^. Goodness-of-fit was assessed by comparing the empirical KS statistic to synthetic datasets drawn from the fitted model; a power law was considered plausible for *p >* 0.1. Likelihood-ratio tests compared power laws to lognormal, exponential, and truncated power-law alternatives.

### Small-world analysis

Small-world structure was quantified for *K* = 3, 5, 10 using the coefficient 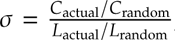, where *C* is mean clustering and *L* mean shortest-path length, compared to an Erdős-Rényi graph *G*(*n, m*) with the same numbers of nodes and edges [110]. Local clustering was computed as *C_i_* = 2*ei* /[*k_i_* (*k_i_* - 1)] and averaged over nodes.

### Assortativity analysis

Degree assortativity *r* was computed as the Pearson correlation between degrees of nodes connected by an edge for *k* = 3, 5, 10. Class-specific assortativity was computed on induced subgraphs, reflecting intra-class connectivity.

### Data analysis

Analyses were performed in Python (3.9.18) using Pandas (1.5.3), NumPy (1.26), SciPy (1.11.3), Polars (0.19.3), Scikit-learn (1.3.2), Scikit-bio (0.5.9), BioPython (1.80), NetworkX (3.1), Seaborn (0.12.2), Matplotlib (3.4.3), Logomaker (0.8), distance (0.1.3), powerlaw (1.5), tqdm (4.66.1) (SI Appendix Table S7), and standard libraries (concurrent.futures, multiprocessing, pathlib, logging, itertools, collections, functools, subprocess, typing, contextlib, statsmodels).

## Supporting Information Appendix (SI)

SI Figures and Tables are appended.

## Data Archival

Code and data are available upon request and will be made publicly available upon peer-reviewed publication.

## Acknowledgments

We thank Klára Hlouchová, Tereza Neuwirthová, Chi Fung Willis Chow, Yuta Nagano, Andreas TiKeau-Mayer, Bharat Ravi Iyengar, and Matthew Merski for discussions and comments. L. A. E. was supported by EMBO (European Molecular Biology Organization) Scientific Exchange Grant 10944. A.T.P. was funded by the Max Planck Gesellschaft (MPG) and ERC (European Research Council) grant (101116284). E.B.-B. acknowledges funding from: HFSP (Human Frontiers of Science Programme, RGP0006/2013 and RGP004/2023); DFG (Deutsche Forschungsgemeinschaft BO-2544/20-1;503272152, BO-2544-22/1;503348080, BO-2544-12/1;298390451); University of Muenster Topical Programme; Volkswagen Foundation (98181), OIST (Okinawa Institute of Science and Technology) Theoretical Sciences Visiting Program 314. Calculations were performed on PALMA II, subsidized by the DFG (INST 211/667-1).

## Supplementary Figures

**Figure 7.**
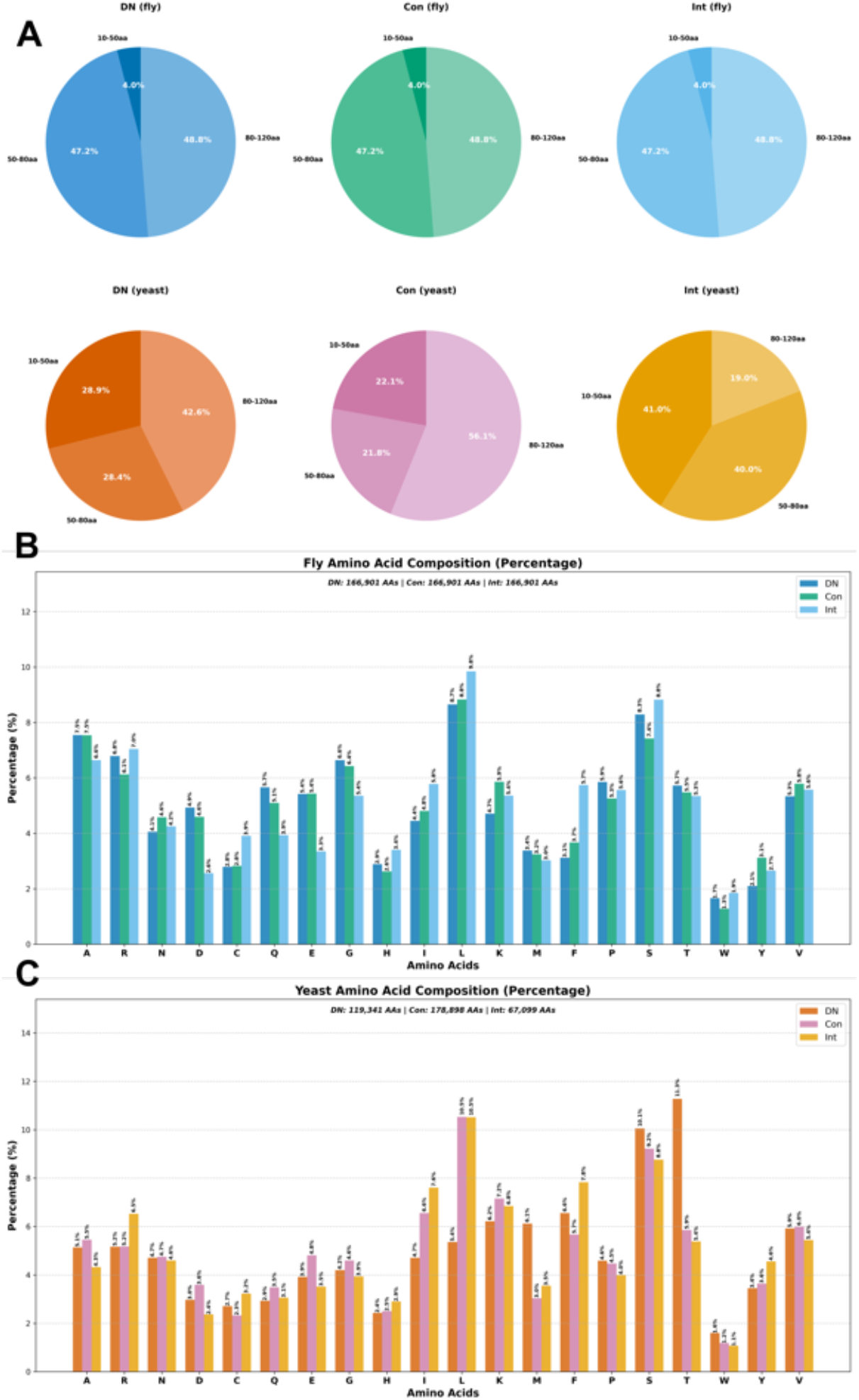
Length distributions and amino-acid composition of fly and yeast sequence sets. **(A)** Protein length-bin distributions (10-50 aa, 50-80 aa, 80-120 aa) for de novo (DN), canonical (Con), and intergenic (Int) sequence sets in fly and yeast (percentages indicated in the pie charts). **(B)** fly amino-acid composition (percent) for DN, Con, and Int sequences, computed from the concatenated sequences in each class (total residues per class shown above the plot). **(C)** yeast amino-acid composition (percent) for DN, Con, and Int sequences, computed analogously (total residues per class shown above the plot).

**Figure 8.**
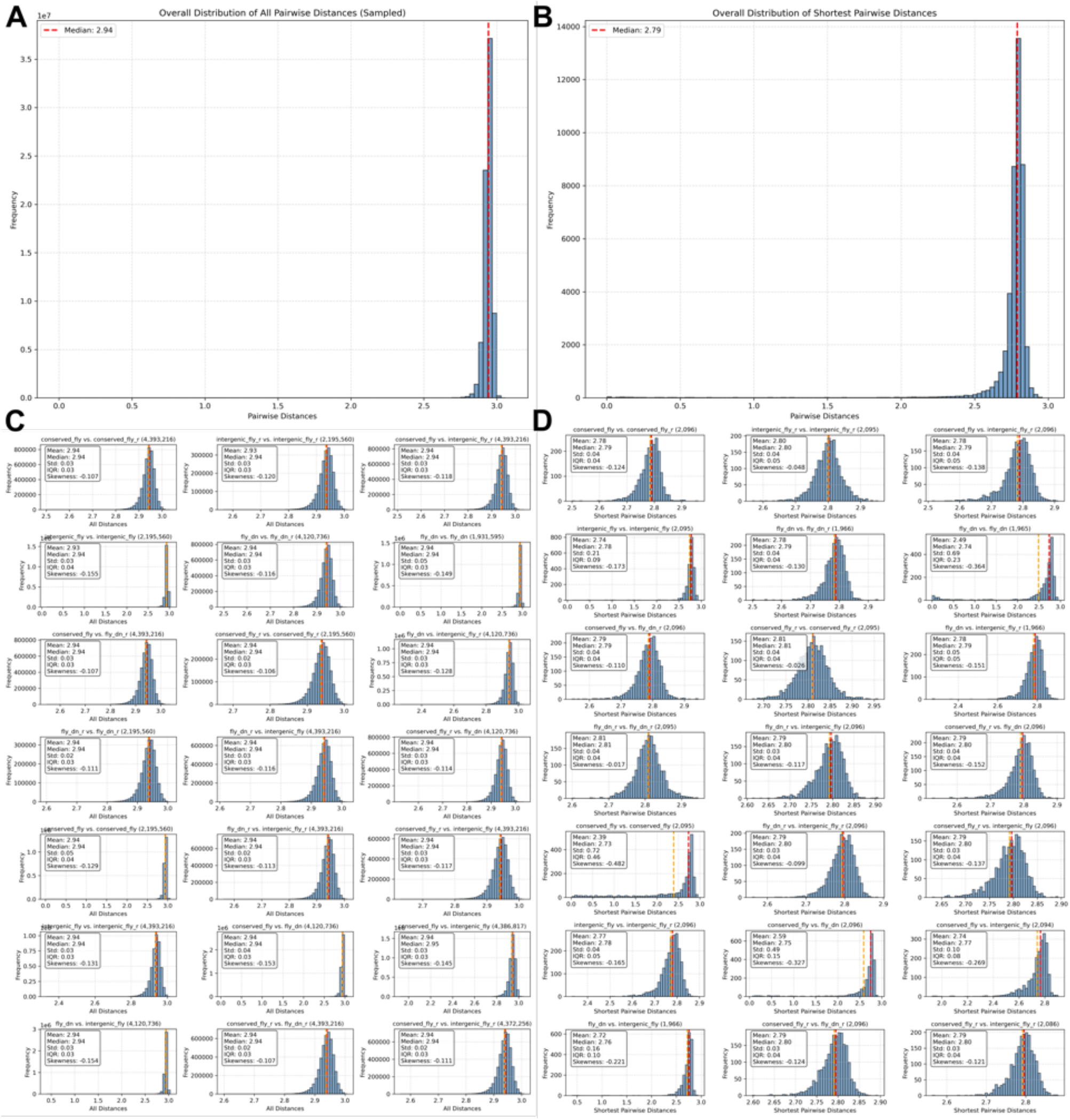
fly distance distributions across sequence classes and randomized controls calculated using SHARK-Dive [74]. **(A)** Overall distribution of sampled *all* pairwise alignment-free distances (as defined in the main text) across the fly data; the dashed red line marks the median. **(B)** Overall distribution of *shortest* pairwise distances (nearest-neighbour distance per sequence); the dashed red line marks the median. **(C)** Pairwise-distance histograms for each class-by-class comparison (DN, Con, Int and their randomized counterparts), with summary statistics reported in each inset; dashed vertical lines indicate median (red) and mean (orange). **(D)** The same set of comparisons as in (C), but using nearest-neighbour (shortest) distances.

**Figure 9.**
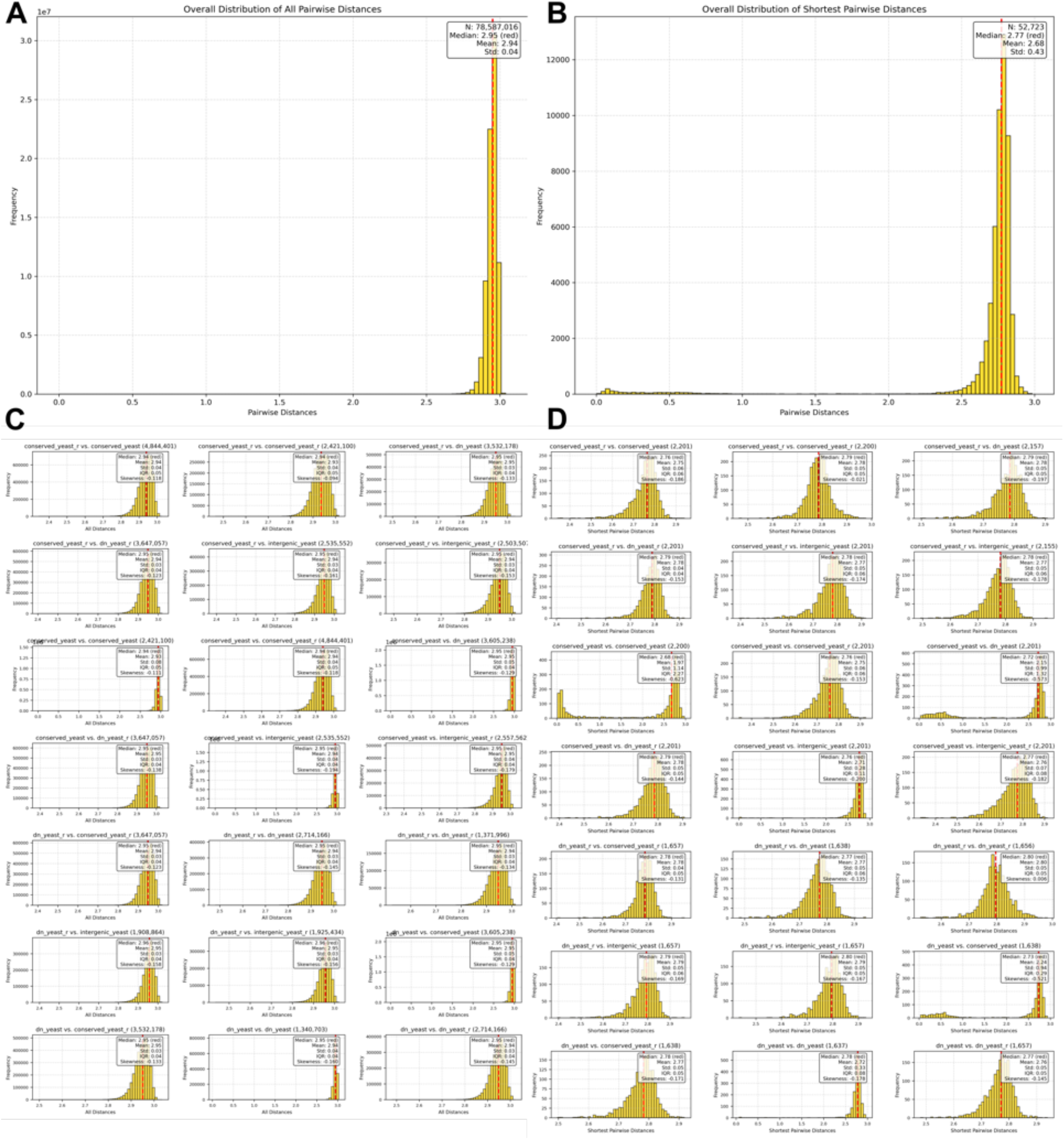
Yeast distance distributions across sequence classes and randomized controls calculated using SHARK-Dive [74]. **(A)** Overall distribution of *all* pairwise alignment-free distances across the yeast data; the dashed red line marks the median (sample size and summary statistics are shown in the panel). **(B)** Overall distribution of nearest-neighbour (shortest) distances; the dashed red line marks the median. **(C)** Pairwise-distance histograms for each class-by-class comparison (DN, Con, Int and randomized counterparts), annotated with per-comparison summary statistics; dashed vertical lines indicate median (red) and mean (orange). **(D)** The same comparisons as in (C), computed for nearest-neighbour (shortest) distances.

**Figure 10.**
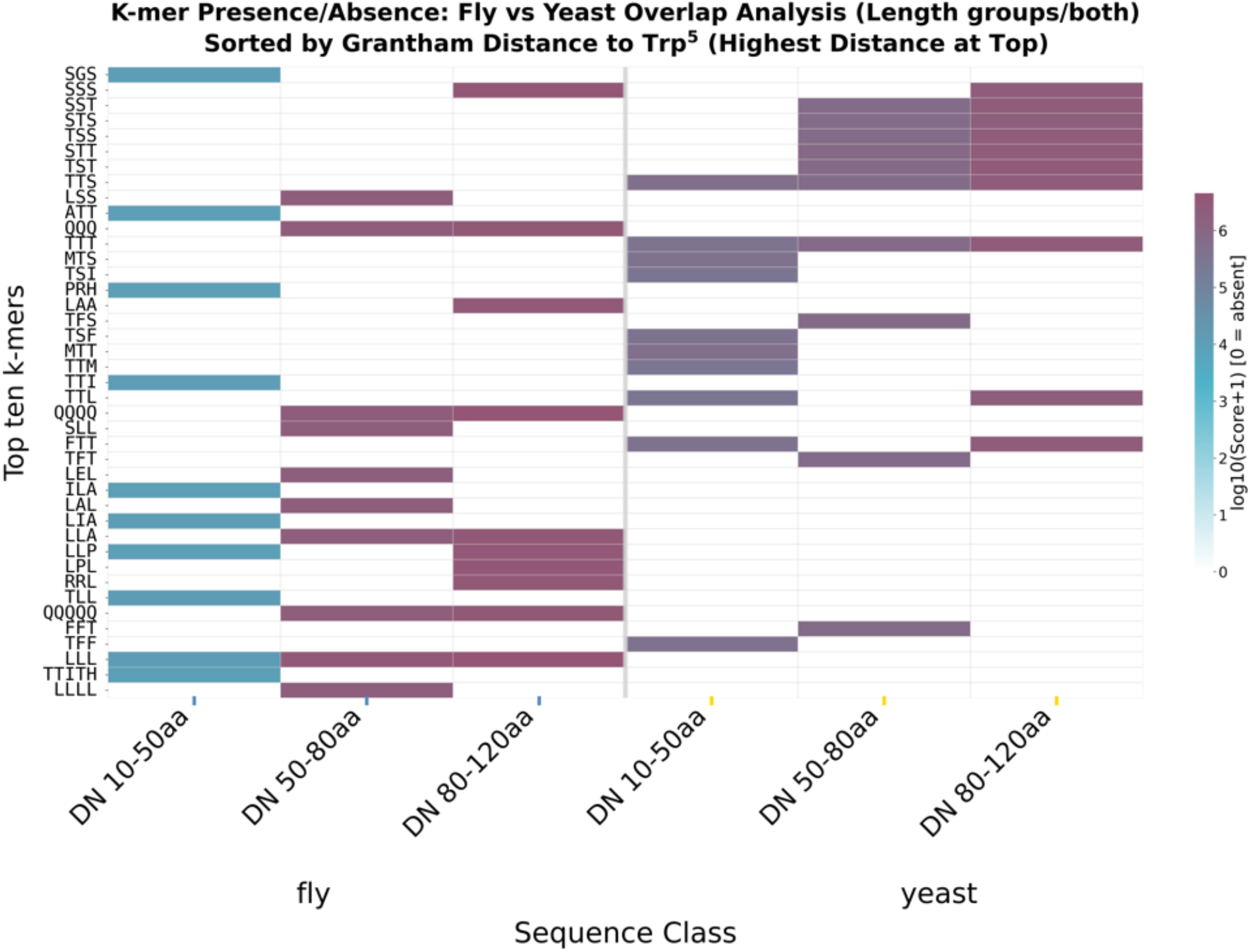
Cross-species *K*-mer enrichment in de novo sequences by length class extracted using SHARK-Capture [75]. Presence/absence heatmap of motifs in de novo (DN) sequences, compared between fly (left block, blue length-class ticks) and yeast (right block, yellow length-class ticks) across the three length bins (10-50 aa, 50-80 aa, 80-120 aa). Colour encodes the enrichment score as log_10_(Score + 1), with 0 indicating an absent *K*-mer. Rows (*K*-mers) are ordered by Grantham distance [76] to tryptophan (Trp), with the highest distance at the top, to highlight compositional divergence from aromatic/bulky residues.

**Figure 11.**
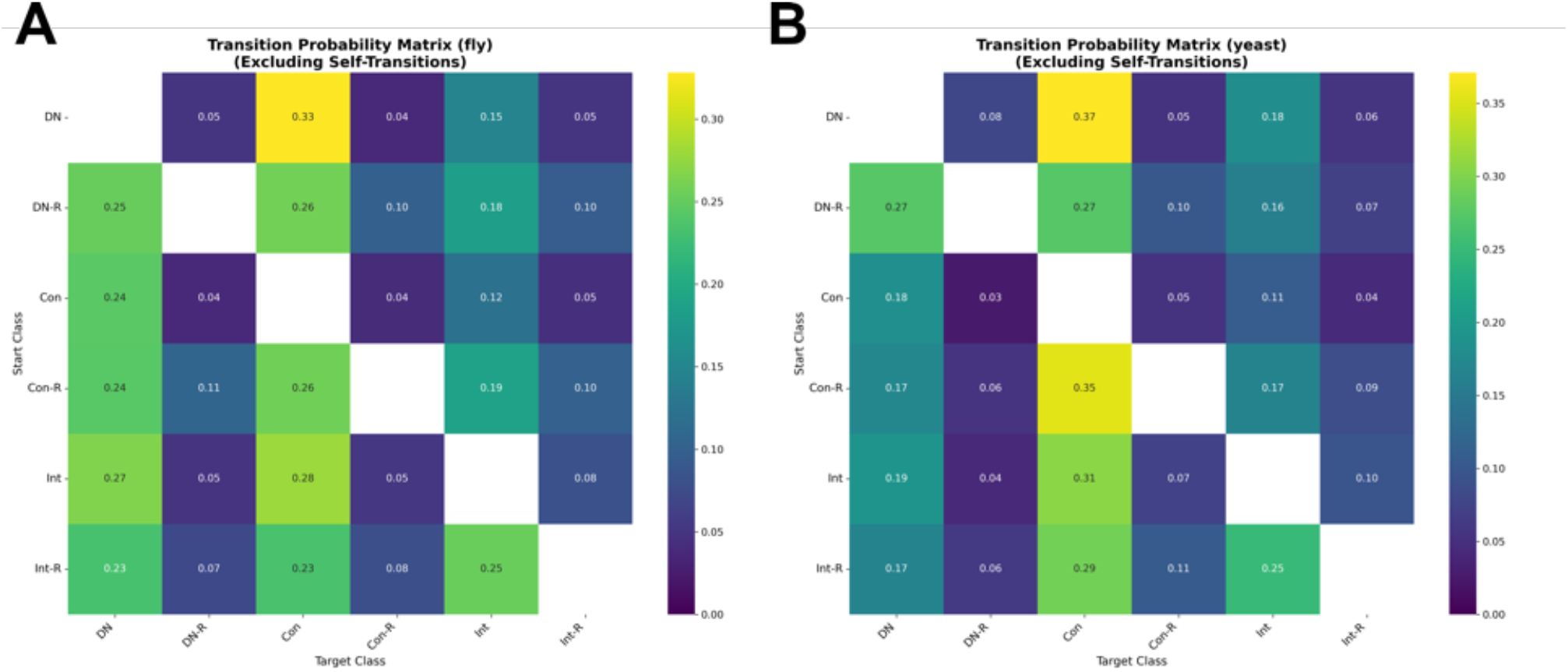
Transition probabilities between sequence classes during greedy traversals. Transition probability matrices (excluding self-transitions) derived from greedy nearest-neighbour traversals in fly (**A**) and yeast (**B**). Each cell gives the probability that a traversal departing from the row class (start class) next reaches the column class (target class), with colour intensity proportional to the transition probability. Classes comprise DN, canonical (Con), intergenic (Int), and their randomized counterparts (DN-R, Con-R, Int-R).

**Figure 12.**
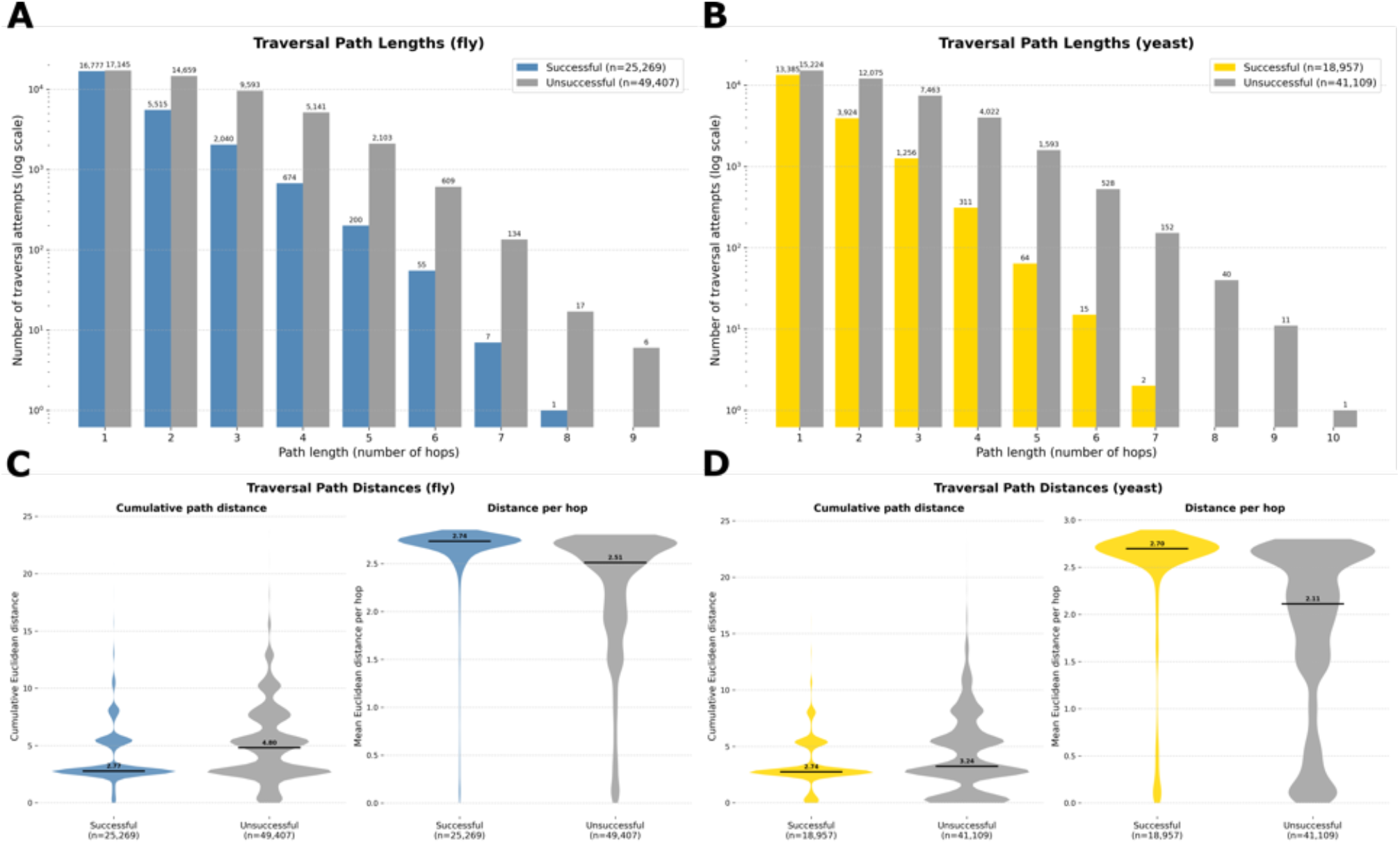
Traversal path lengths and path distances for successful and unsuccessful traversals. **(A-B)** Distribution of traversal path lengths (number of hops) for successful versus unsuccessful traversal attempts in fly (**A**) and yeast (**B**); bar heights give the number of traversal attempts on a logarithmic scale (sample sizes indicated in-panel). **(C-D)** Traversal path distances for fly (**C**) and yeast (**D**), comparing successful and unsuccessful traversals. Left violins show the cumulative Euclidean path distance per traversal; right violins show the mean per-hop Euclidean distance. Overlaid markers indicate central tendency.

**Figure 13.**
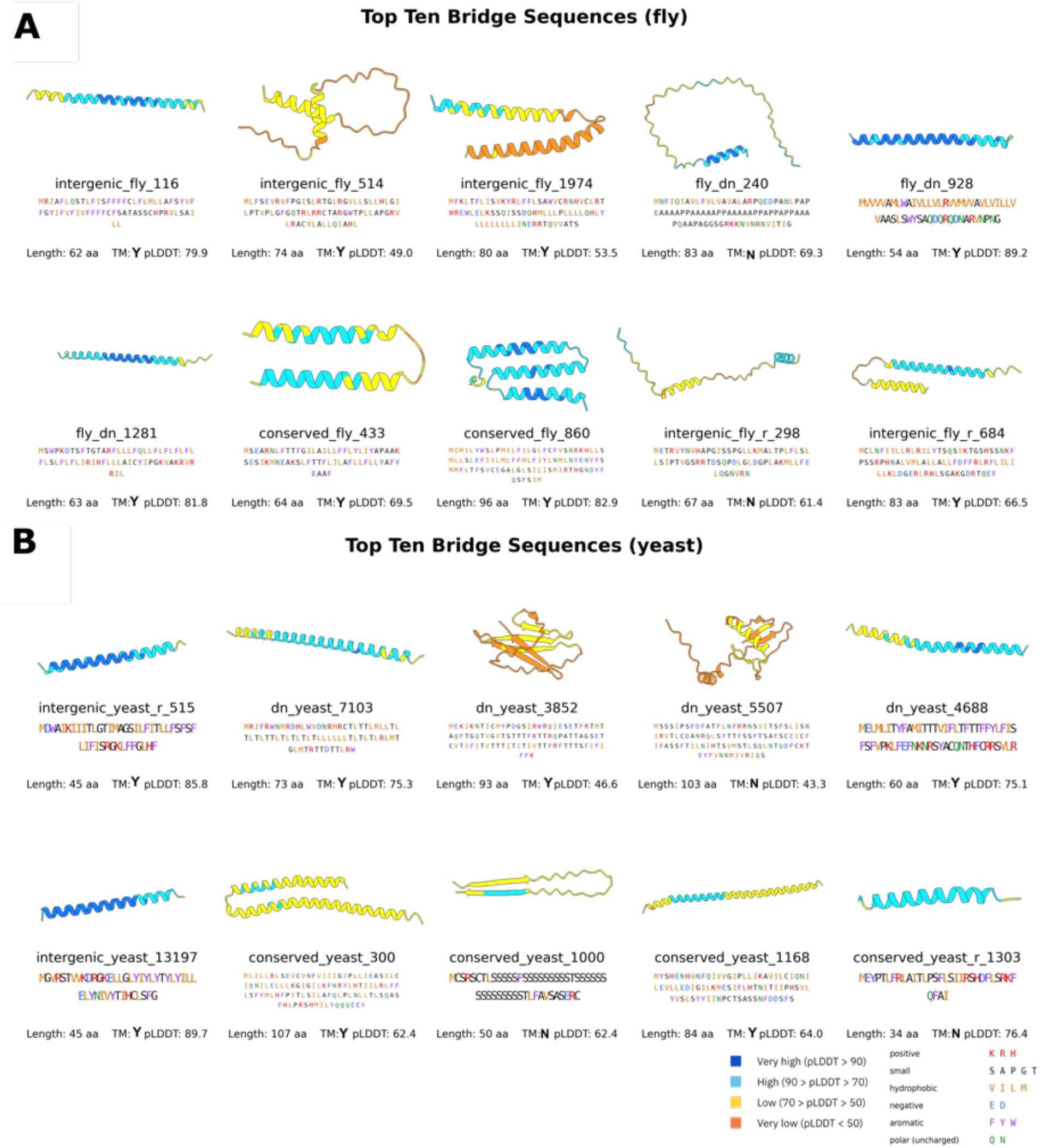
Top-ten bridge sequences and their predicted structures. AlphaFold3 structure predictions for the top ten bridge sequences, ranked by raw intermediate-use counts across successful traversals, in fly (**A**) and yeast (**B**). Each panel reports the sequence identifier and class membership, the primary sequence, length (aa), predicted transmembrane status (TM: Y/N), and mean pLDDT. Cartoons are coloured by per-residue pLDDT confidence ([105]) (very high *>* 90; high 70-90; low 50-70; very low *<* 50); sequence text is coloured by residue physicochemical class (positive, small, hydrophobic, negative, aromatic, polar/uncharged) as indicated in the legend. Transmembrane enrichment statistics for these sets are reported in SI Appendix Fig. S10. Structures visualized in UCSF ChimeraX ([111]).

**Figure 14.**
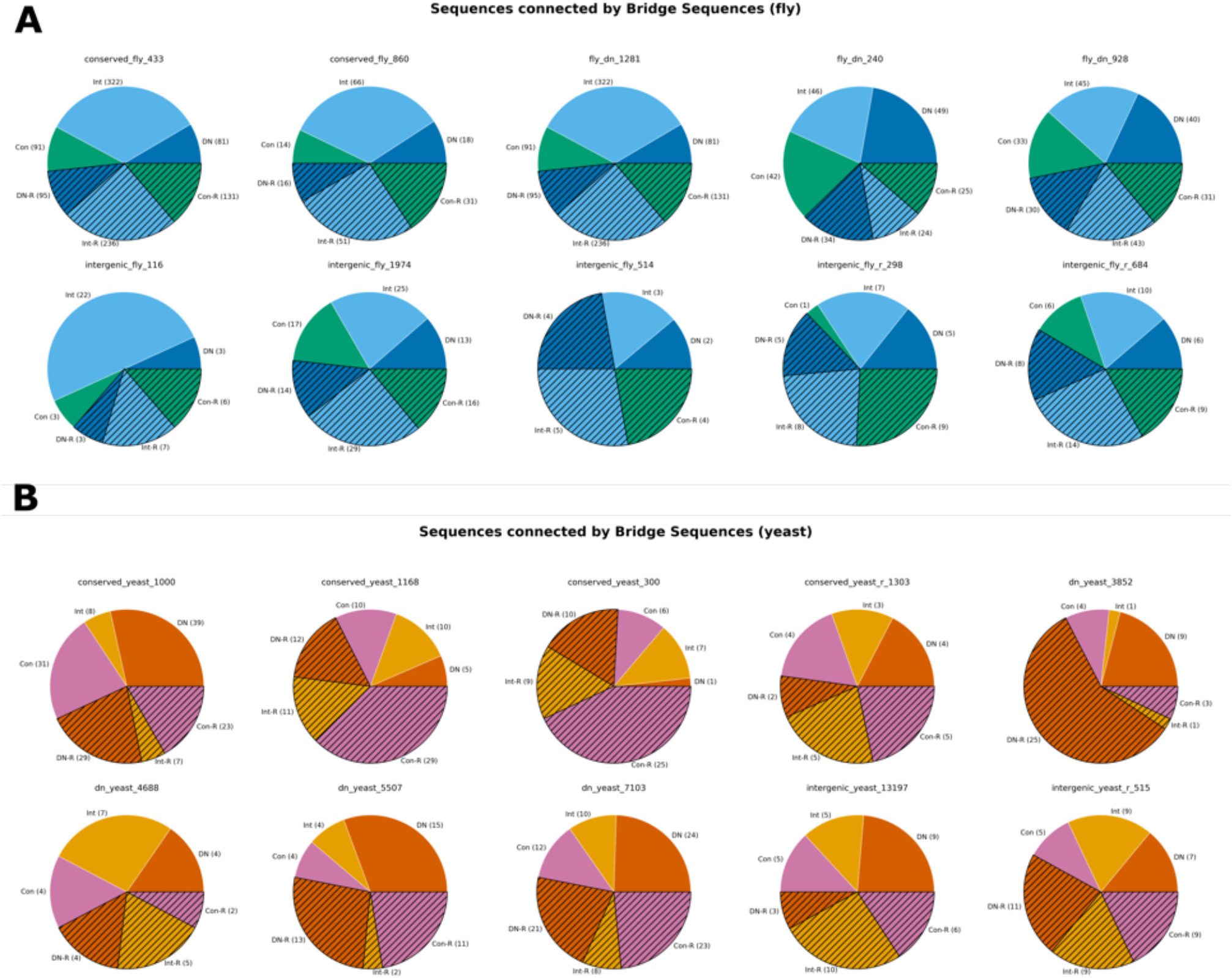
Class composition of bridge-connected neighbourhoods. Pie charts showing the class composition of sequences connected by each of the top ten bridge sequences in fly (**A**) and yeast (**B**). For each bridge, the chart gives the proportion of connected neighbours belonging to each class (DN, Con, Int, and their randomized counterparts DN-R, Con-R, Int-R), with absolute counts indicated. Bridge-sequence identifiers and class membership are labelled above each chart.

**Figure 15.**
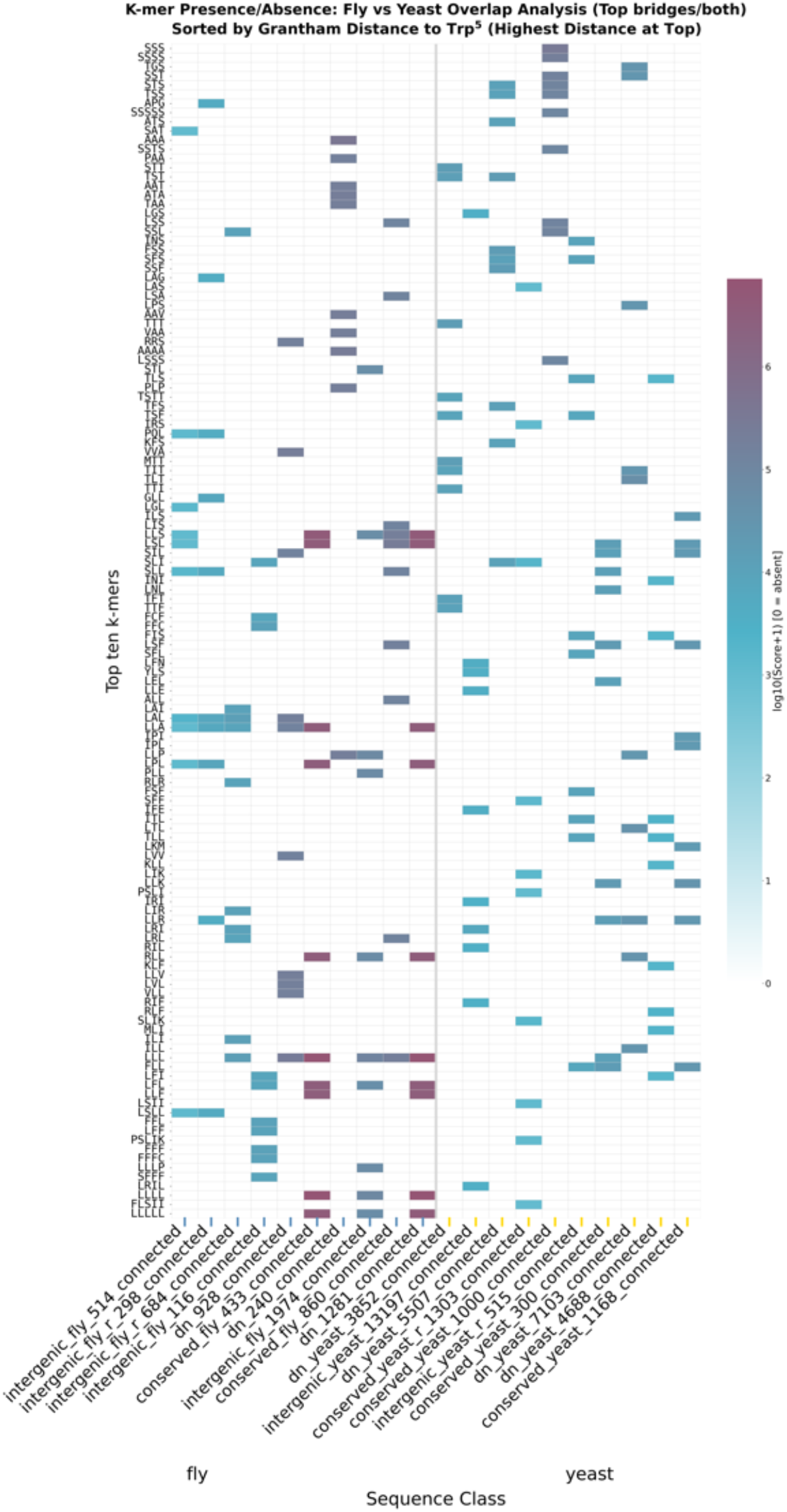
Cross-species *K*-mer enrichment among bridge-connected sequences. Presence/absence heatmap of enriched motifs (SHARK-Capture [75]) extracted from sequences connected to the top bridge sequences, compared between fly (blue class ticks) and yeast (yellow class ticks) across sequence classes and their randomized controls. Colour encodes the enrichment score as log_10_(Score + 1), with 0 indicating an absent *K*-mer. Rows (*K*-mers) are ordered by Grantham distance [76] to tryptophan (Trp), with the highest distance at the top.

**Figure 16.**
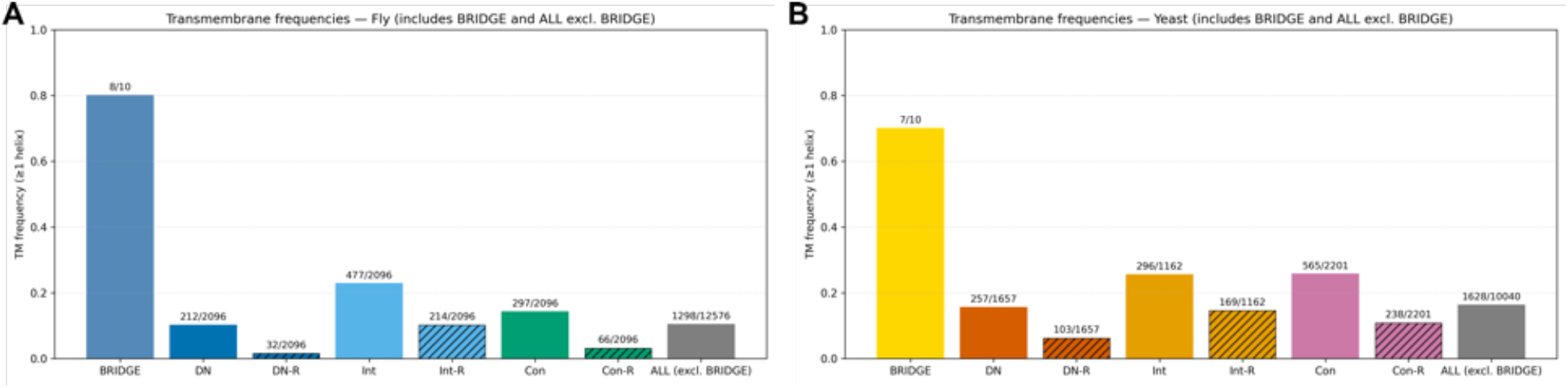
Transmembrane (TM) enrichment across sequence classes predicted using Phobius ([108]) Frequency of predicted transmembrane signal (at least one TM helix) across sequence sets in fly (**A**) and yeast (**B**), including the BRIDGE set and all sequences excluding BRIDGE (ALL excl. BRIDGE), as well as DN, canonical (Con), intergenic (Int), and their randomized counterparts (DN-R, Con-R, Int-R). Counts (TM-positive / total) are shown above each bar. Bridge sequences are strongly enriched for TM signal relative to species-wide backgrounds: in fly, 8/10 (95% CI = 0.49-0.94) contain at least one TM helix (Fisher’s exact, *q* = 9.8 x10^-7^); in yeast, 7/10 (95% CI = 0.40-0.89) are TM-positive (*q* = 2.3 x 10^-4^). Confidence intervals are 95% Wilson intervals and significance is from FDR-corrected Fisher’s exact tests.

**Figure 17.**
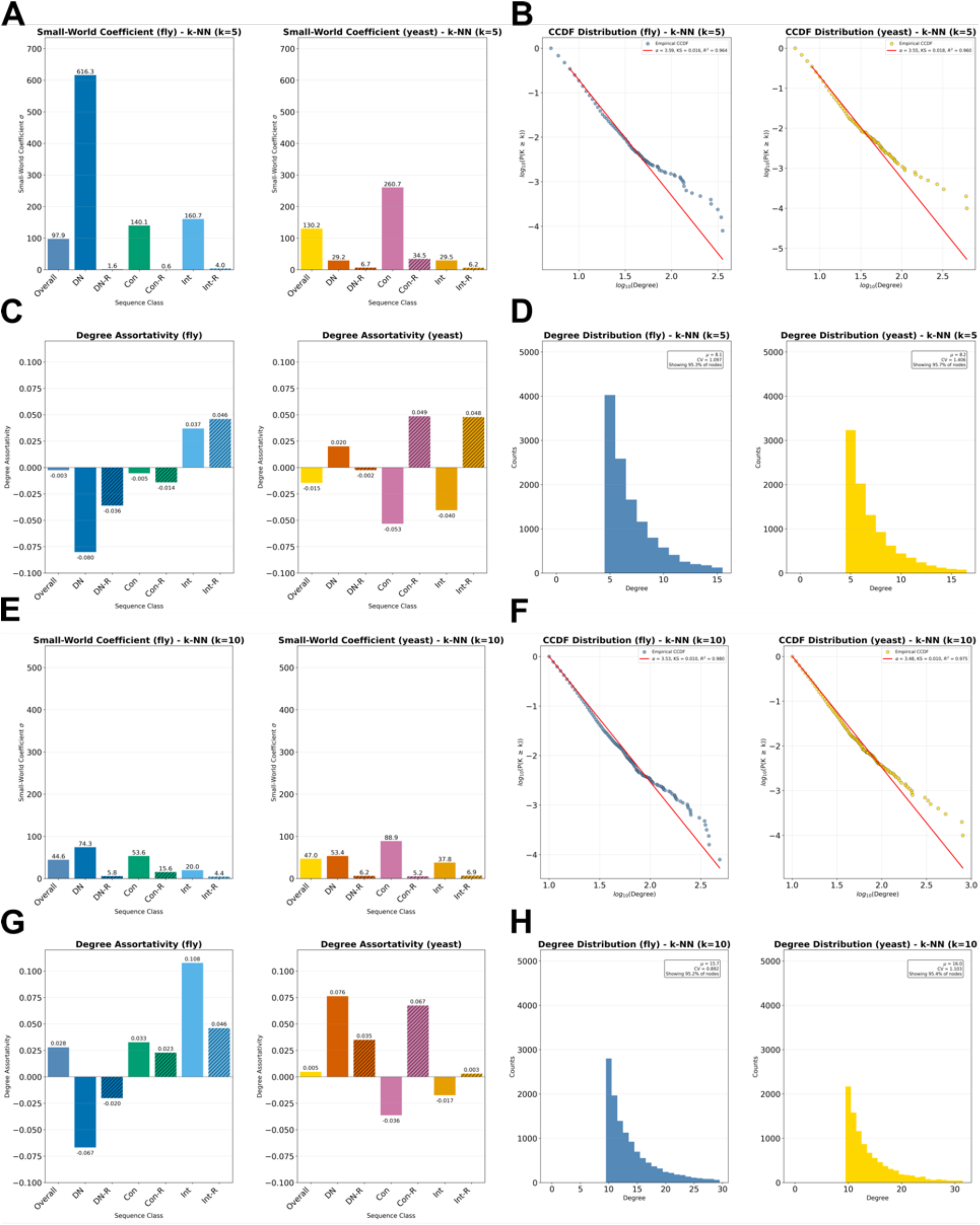
Graph topology of fly and yeast k-NN networks. Network summary statistics computed on *k*-nearest-neighbour (k-NN) graphs for fly and yeast, shown for *K* = 5 (**A-D**) and *K* = 10 (**E-H**). **(A, E)** Small-world coefficient (σ) for the overall graph and for each sequence class (including randomized controls). **(B, F)** Degree-distribution complementary cumulative distribution functions (CCDFs) ([109]) on log-log axes, with a power-law fit overlaid (as indicated in-panel). **(C, G)** Degree assortativity for overall graphs and per class. **(D, H)** Degree distributions (histograms) for fly and yeast k-NN graphs.

**Figure 18.**
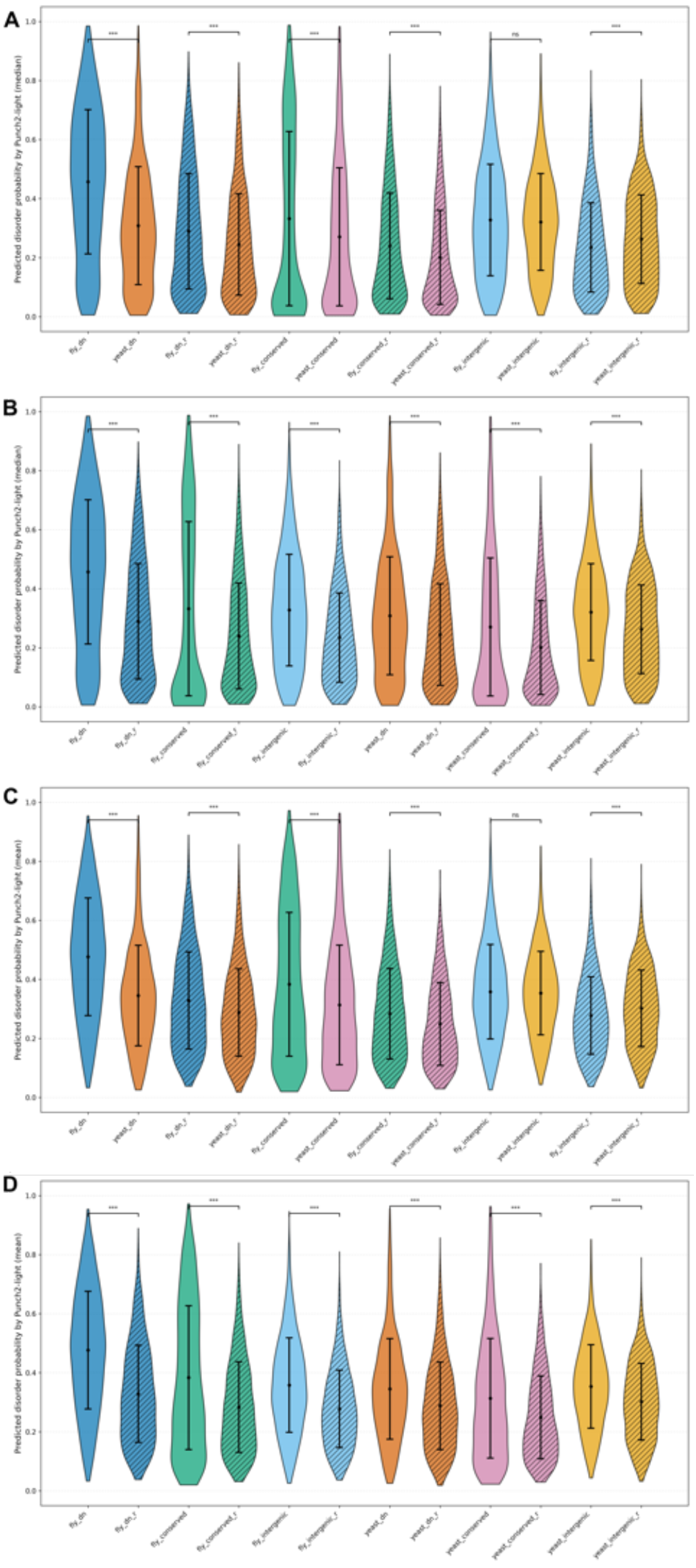
Predicted intrinsic disorder across classes, species, and randomized controls using Punch2-light ([107]) Violin plots show per-sequence predicted disorder probabilities from punch2-light (central tendency indicated by the overlaid point and error bars). **(A)** Median disorder probability comparing fly vs. yeast for DN, canonical, and intergenic sequences (and their randomized controls; significance annotations above brackets). **(B)** Median disorder probability comparing original vs. randomized sequences within fly and within yeast. **(C)** Same as (A), but using mean disorder probability per sequence. **(D)** Same as (B), but using mean disorder probability per sequence.

## Supplementary Tables

**Table 1.**
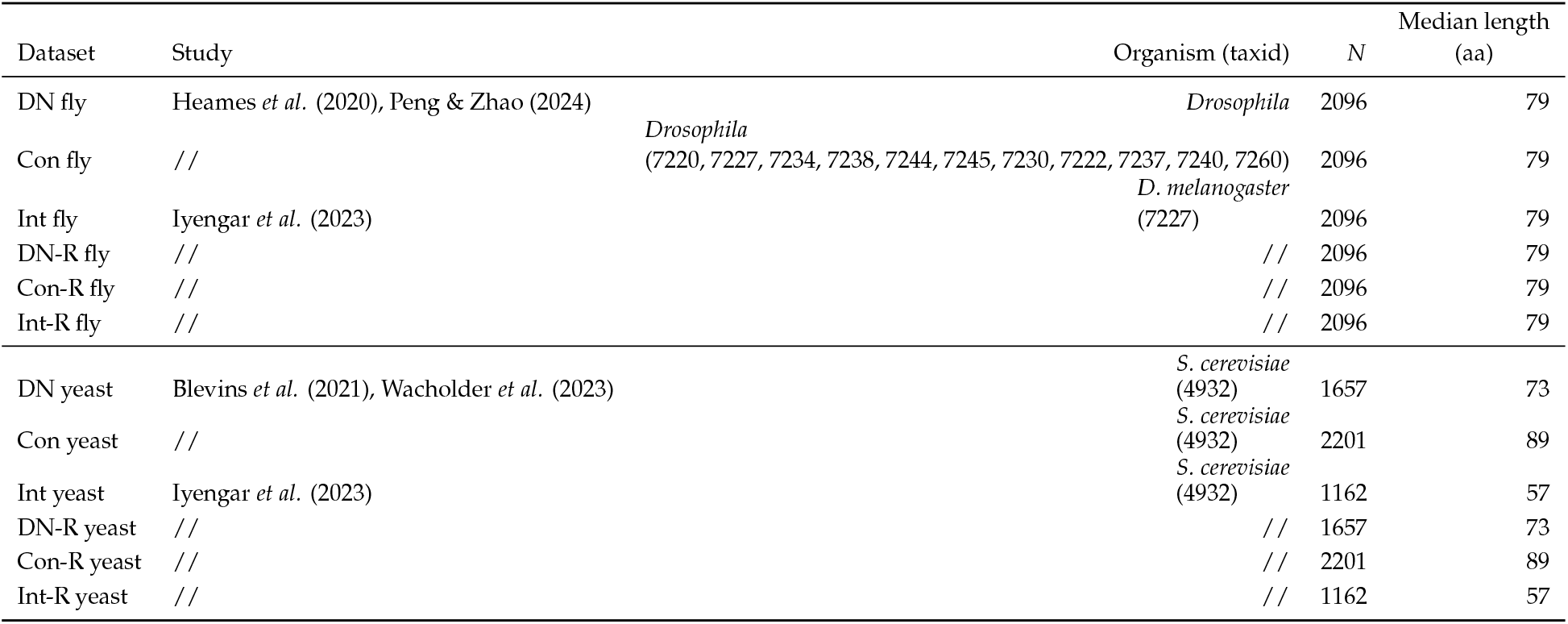
Datasets used in this study (10-120 aa, non-redundant, canonical amino acids). During analysis, residual duplicates (fly: 20; yeast: 5) were removed because they produced zero pairwise distances and were found to be identical and escaped initial filtering using SeqKit ([112]).

**Table 2.** Original Grantham distance matrix for the 20 standard amino acids [. **76].**

|  | S | R | L | P | T | A | V | G | I | F | Y | C | H | Q | N | K | D | E | M | W |
| --- | --- | --- | --- | --- | --- | --- | --- | --- | --- | --- | --- | --- | --- | --- | --- | --- | --- | --- | --- | --- |
| S | 0 | 110 | 145 | 74 | 58 | 99 | 124 | 56 | 142 | 155 | 144 | 112 | 89 | 68 | 46 | 121 | 65 | 80 | 135 | 177 |
| R | 110 | 0 | 102 | 103 | 71 | 112 | 96 | 125 | 97 | 97 | 77 | 180 | 29 | 43 | 86 | 26 | 96 | 54 | 91 | 101 |
| L | 145 | 102 | 0 | 98 | 92 | 96 | 32 | 138 | 5 | 22 | 36 | 198 | 99 | 113 | 153 | 107 | 172 | 138 | 15 | 61 |
| P | 74 | 103 | 98 | 0 | 38 | 27 | 68 | 42 | 95 | 114 | 110 | 169 | 77 | 76 | 91 | 103 | 108 | 93 | 87 | 147 |
| T | 58 | 71 | 92 | 38 | 0 | 58 | 69 | 59 | 89 | 103 | 92 | 149 | 47 | 42 | 65 | 78 | 85 | 65 | 81 | 128 |
| A | 99 | 112 | 96 | 27 | 58 | 0 | 64 | 60 | 94 | 113 | 112 | 195 | 86 | 91 | 111 | 106 | 126 | 107 | 84 | 148 |
| V | 124 | 96 | 32 | 68 | 69 | 64 | 0 | 109 | 29 | 50 | 55 | 192 | 84 | 96 | 133 | 97 | 152 | 121 | 21 | 88 |
| G | 56 | 125 | 138 | 42 | 59 | 60 | 109 | 0 | 135 | 153 | 147 | 159 | 98 | 87 | 80 | 127 | 94 | 98 | 127 | 184 |
| I | 142 | 97 | 5 | 95 | 89 | 94 | 29 | 135 | 0 | 21 | 33 | 198 | 94 | 109 | 149 | 102 | 168 | 134 | 10 | 61 |
| F | 155 | 97 | 22 | 114 | 103 | 113 | 50 | 153 | 21 | 0 | 22 | 205 | 100 | 116 | 158 | 102 | 177 | 140 | 28 | 40 |
| Y | 144 | 77 | 36 | 110 | 92 | 112 | 55 | 147 | 33 | 22 | 0 | 194 | 83 | 99 | 143 | 85 | 160 | 122 | 36 | 37 |
| C | 112 | 180 | 198 | 169 | 149 | 195 | 192 | 159 | 198 | 205 | 194 | 0 | 174 | 154 | 139 | 202 | 154 | 170 | 196 | 215 |
| H | 89 | 29 | 99 | 77 | 47 | 86 | 84 | 98 | 94 | 100 | 83 | 174 | 0 | 24 | 68 | 32 | 81 | 40 | 87 | 115 |
| Q | 68 | 43 | 113 | 76 | 42 | 91 | 96 | 87 | 109 | 116 | 99 | 154 | 24 | 0 | 46 | 53 | 61 | 29 | 101 | 130 |
| N | 46 | 86 | 153 | 91 | 65 | 111 | 133 | 80 | 149 | 158 | 143 | 139 | 68 | 46 | 0 | 94 | 23 | 42 | 142 | 174 |
| K | 121 | 26 | 107 | 103 | 78 | 106 | 97 | 127 | 102 | 102 | 85 | 202 | 32 | 53 | 94 | 0 | 101 | 56 | 95 | 110 |
| D | 65 | 96 | 172 | 108 | 85 | 126 | 152 | 94 | 168 | 177 | 160 | 154 | 81 | 61 | 23 | 101 | 0 | 45 | 160 | 181 |
| E | 80 | 54 | 138 | 93 | 65 | 107 | 121 | 98 | 134 | 140 | 122 | 170 | 40 | 29 | 42 | 56 | 45 | 0 | 126 | 152 |
| M | 135 | 91 | 15 | 87 | 81 | 84 | 21 | 127 | 10 | 28 | 36 | 196 | 87 | 101 | 142 | 95 | 160 | 126 | 0 | 67 |
| W | 177 | 101 | 61 | 147 | 128 | 148 | 88 | 184 | 61 | 40 | 37 | 215 | 115 | 130 | 174 | 110 | 181 | 152 | 67 | 0 |

**Table 3.** Wasserstein-1 (*W*_1_) ([100]), k-sample Anderson-Darling statistic (AD) ([101]), and squared Maximum Mean Discrepancy (MMD^2^) ([102]) for every comparison in fly and yeast, overall and within three sequence-length bins. *p*_BH_ is the Benjamini-Hochberg-adjusted permutation *p*-value of MMD^2^ within each bucket across both species ([103]); reject indicates rejection of the null of identical distributions at BH-adjusted *p <* 0.05 (✓) or failure to reject (×).

| Species | Bucket | Comparison | $W_1$ | AD | MMD<sup>2</sup> | $p_{BH}$ | reject |
| --- | --- | --- | --- | --- | --- | --- | --- |
|  | 50-80 aa |  | 0.0023 | 150 | 2.04e-06 | 0.002 | ✓ |
|  | 80-120 aa |  | 0.0020 | 48.9 | 2.53e-06 | 0.002 | ✓ |
|  | overall | Con vs Con to Con vs Con-R | 0.0048 | 3,328 | 1.32e-06 | 0.002 | ✓ |
|  | 10-50 aa |  | 0.0064 | 276 | 1.13e-05 | 0.003 | ✓ |
|  | 50-80 aa |  | 0.0079 | 543 | 6.92e-06 | 0.002 | ✓ |
|  | 80-120 aa |  | 0.0084 | 5,847 | 3.02e-06 | 0.002 | ✓ |
|  | overall | Con vs DN to Con vs DN-R | 0.0026 | 1,681 | 5.87e-07 | 0.002 | ✓ |
|  | 10-50 aa |  | 0.0017 | 83.1 | 1.19e-07 | 0.031 | ✓ |
|  | 50-80 aa |  | 0.0035 | 213 | 3.20e-06 | 0.002 | ✓ |
|  | 80-120 aa |  | 0.0048 | 2,077 | 9.85e-07 | 0.002 | ✓ |
|  | overall | Con vs Int to Con vs Int-R | 0.0026 | 1,364 | 6.26e-06 | 0.002 | ✓ |
|  | 10-50 aa |  | 0.0035 | 142 | 1.16e-05 | 0.003 | ✓ |
|  | 50-80 aa |  | 0.0024 | 130 | 3.16e-06 | 0.002 | ✓ |
|  | 80-120 aa |  | 0.0030 | 210 | 5.61e-06 | 0.002 | ✓ |
|  | overall | Int vs Int to Int vs Int-R | 0.0041 | 1,445 | 1.83e-05 | 0.002 | ✓ |
|  | 10-50 aa |  | 0.0046 | 234 | 2.33e-05 | 0.003 | ✓ |
|  | 50-80 aa |  | 0.0045 | 340 | 1.99e-05 | 0.002 | ✓ |
|  | 80-120 aa |  | 0.0051 | 81.7 | 2.35e-05 | 0.002 | ✓ |
|  | overall | Int vs DN to Int vs DN-R | 0.0022 | 1,445 | 1.45e-06 | 0.002 | ✓ |
|  | 10-50 aa |  | 0.0019 | 98.4 | 2.92e-06 | 0.003 | ✓ |
|  | 50-80 aa |  | 0.0040 | 561 | 1.14e-05 | 0.002 | ✓ |
|  | 80-120 aa |  | 0.0038 | 668 | 3.51e-07 | 0.002 | ✓ |
|  | overall | Int vs Con to Int vs Con-R | 0.0038 | 3,725 | 3.31e-06 | 0.002 | ✓ |
|  | 10-50 aa |  | 0.0046 | 288 | 2.13e-05 | 0.003 | ✓ |
|  | 50-80 aa |  | 0.0036 | 713 | 2.23e-07 | 0.004 | ✓ |
|  | 80-120 aa |  | 0.0066 | 1,815 | 8.24e-06 | 0.002 | ✓ |

**Table 4.** Original top 10 bridge sequences with statistical validation, Fly.

| Sequence<br>TM | BridgeScore | Obs_N1 | Mean_N1 | SD_N1 | Z_N1 | P_N1 | FDR_Q_N1 | FoldEnr_N1 | Obs_N2 | Mean_N2 | SD_N2 | Z_N2 | P_N2 | FDR_Q_N2 | FoldEnr_N2 | FisherP_N1 | FisherQ_N1 | SP |
| --- | --- | --- | --- | --- | --- | --- | --- | --- | --- | --- | --- | --- | --- | --- | --- | --- | --- | --- |
| fly_dn_1281<br>1 | 553 | 553.0 | 102.11 | 83.78 | 5.38 | 0.005 | 1.000 | 5.42 | 553.0 | 241.88 | 15.36 | 20.25 | 0.005 | 1.000 | 2.29 | 0.350 | 6.23e-04 | no |
| canonical_fly_860<br>3 | 174 | 174.0 | 32.10 | 27.90 | 5.09 | 0.005 | 1.000 | 5.42 | 174.0 | 89.22 | 9.50 | 8.93 | 0.005 | 1.000 | 1.95 | 0.350 | 6.23e-04 | no |
| intergenic_fly_197<br>1 | 153 | 153.0 | 32.42 | 20.68 | 5.83 | 0.005 | 1.000 | 4.72 | 153.0 | 85.91 | 9.08 | 7.39 | 0.005 | 1.000 | 1.78 | 3.51e-04 | 6.23e-04 | yes |
| fly_dn_240<br>0 | 134 | 134.0 | 47.65 | 37.51 | 2.30 | 0.025 | 0.848 | 2.81 | 134.0 | 86.84 | 9.44 | 4.99 | 0.005 | 1.000 | 1.54 | 0.674 | 6.23e-04 | yes |
| canonical_fly_433<br>2 | 83 | 83.0 | 28.17 | 21.65 | 2.53 | 0.020 | 0.848 | 2.95 | 83.0 | 91.71 | 9.52 | -0.91 | 0.841 | 1.000 | 0.91 | 0.633 | 6.23e-04 | no |
| fly_dn_928<br>1 | 81 | 81.0 | 34.17 | 25.00 | 1.87 | 0.070 | 0.848 | 2.37 | 81.0 | 61.12 | 7.60 | 2.62 | 0.010 | 1.000 | 1.33 | 0.697 | 6.23e-04 | no |
| intergenic_fly_116<br>1 | 78 | 78.0 | 19.18 | 13.02 | 4.52 | 0.005 | 1.000 | 4.07 | 78.0 | 42.82 | 6.04 | 5.82 | 0.005 | 1.000 | 1.82 | 0.215 | 6.23e-04 | no |
| intergenic_fly_r_68<br>1 | 70 | 70.0 | 17.52 | 14.18 | 3.70 | 0.015 | 0.848 | 3.99 | 70.0 | 65.64 | 8.68 | 0.50 | 0.338 | 1.000 | 1.07 | 0.002 | 6.23e-04 | no |
| intergenic_fly_r_29<br>0 | 50 | 50.0 | 2.28 | 3.05 | 15.65 | 0.005 | 0.885 | 21.93 | 50.0 | 12.61 | 3.70 | 10.09 | 0.005 | 1.000 | 3.97 | 7.21e-06 | 6.23e-04 | no |
| intergenic_fly_514<br>1 | 44 | 44.0 | 3.07 | 3.03 | 13.53 | 0.005 | 1.000 | 14.33 | 44.0 | 12.85 | 3.41 | 9.14 | 0.005 | 1.000 | 3.43 | 5.01e-08 | 6.23e-04 | no |

**Table 5.**
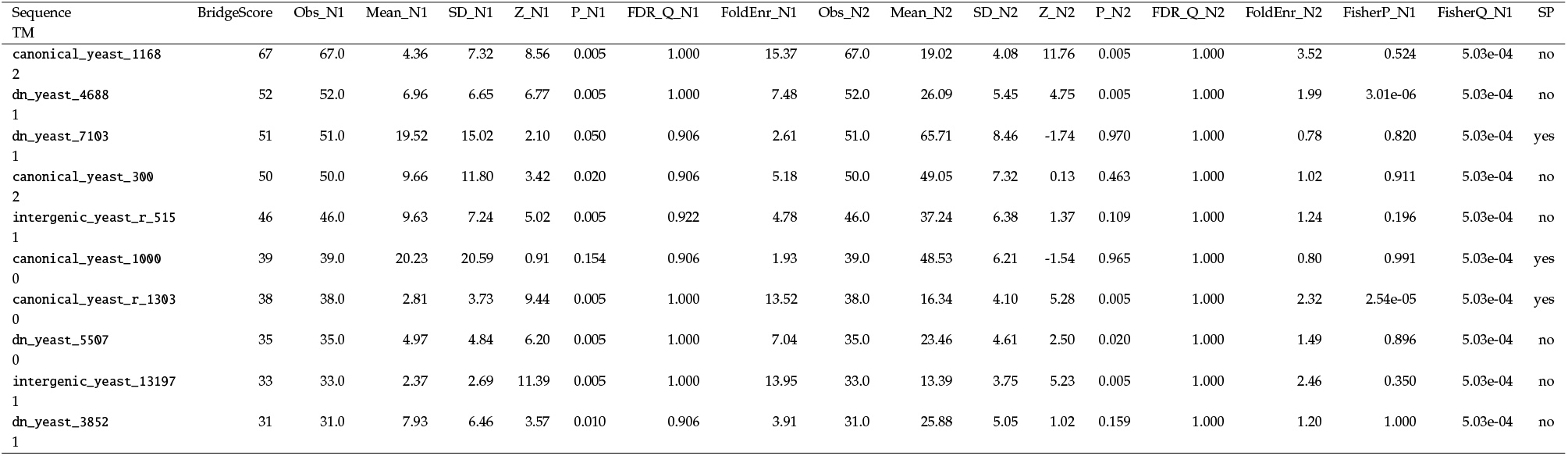
Original top 10 bridge sequences with statistical validation, Yeast.

**Table 6.** TM bridge correlation results.

| Species | Class | N_total | N_with_TM | N_without_TM | Spearman_rho | Spearman_p_adj | Spearman_sig | Median_BS_with_TM | Median_BS_without_TM | MannWhitney_p_adj | MannWhitney_sig |
| --- | --- | --- | --- | --- | --- | --- | --- | --- | --- | --- | --- |
| Fly | DN | 2096 | 212 | 1884 | 0.0591 | 1.03e-02 | * | 0.0 | 0.0 | 1.12e-02 | * |
| Fly | Con | 2096 | 297 | 1799 | 0.0859 | 2.11e-04 | *** | 0.0 | 0.0 | 2.47e-04 | *** |
| Fly | Int | 2096 | 477 | 1619 | 0.1804 | 1.02e-15 | *** | 0.0 | 0.0 | 2.17e-14 | *** |
| Fly | DN-R | 2096 | 32 | 2064 | -0.0046 | 8.32e-01 | ns | 0.0 | 0.0 | 8.26e-01 | ns |
| Fly | Con-R | 2096 | 66 | 2030 | 0.0574 | 1.14e-02 | * | 0.0 | 0.0 | 1.12e-02 | * |
| Fly | Int-R | 2096 | 214 | 1882 | 0.0751 | 1.10e-03 | ** | 0.0 | 0.0 | 1.30e-03 | ** |
| Yeast | DN | 1657 | 257 | 1400 | 0.153 | 2.28e-09 | *** | 0.0 | 0.0 | 2.32e-09 | *** |
| Yeast | Con | 2201 | 565 | 1636 | 0.0499 | 2.31e-02 | * | 0.0 | 0.0 | 4.08e-02 | * |
| Yeast | Int | 1162 | 296 | 866 | 0.1 | 1.10e-03 | ** | 0.0 | 0.0 | 1.87e-03 | ** |
| Yeast | DN-R | 1657 | 103 | 1554 | 0.0357 | 1.60e-01 | ns | 0.0 | 0.0 | 1.58e-01 | ns |
| Yeast | Con-R | 2201 | 238 | 1963 | 0.0835 | 2.11e-04 | *** | 0.0 | 0.0 | 2.47e-04 | *** |
| Yeast | Int-R | 1162 | 169 | 993 | 0.124 | 9.02e-05 | *** | 0.0 | 0.0 | 1.22e-04 | *** |

**Table 7.**
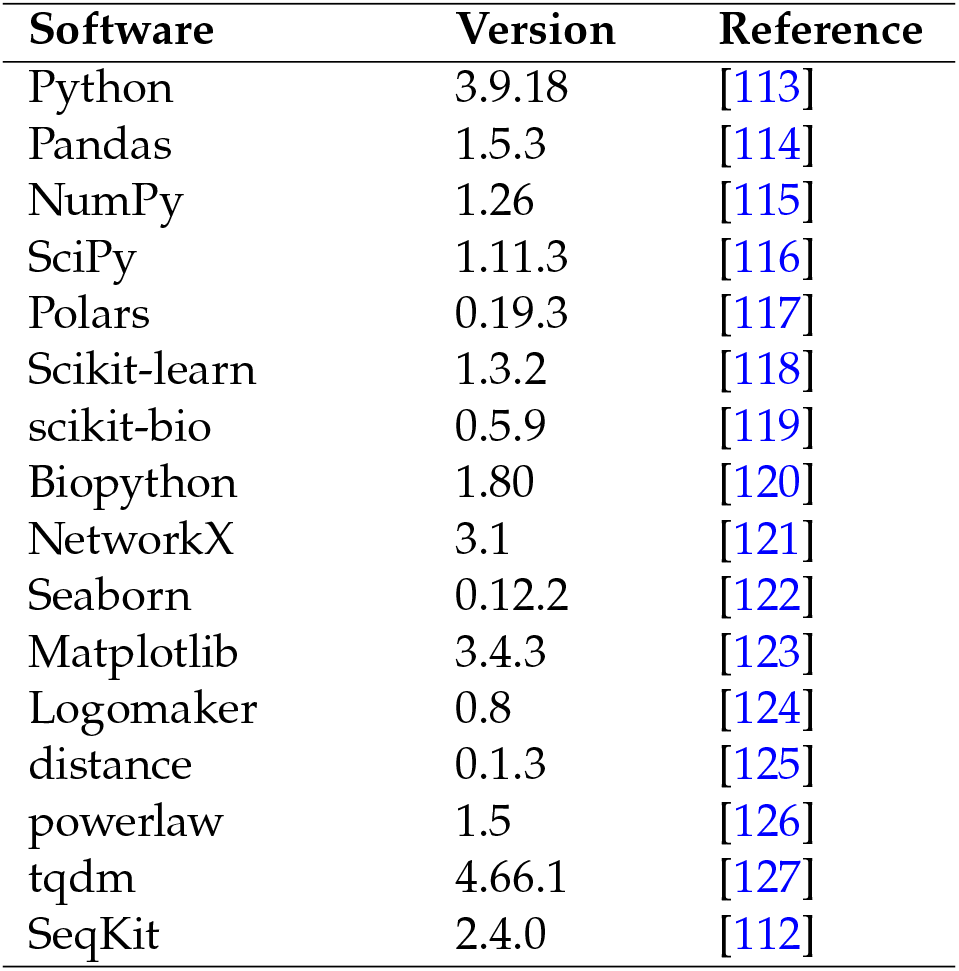
Software libraries used for the analyses.

